# Widespread 3D genome reorganization precedes programmed DNA rearrangement in *Oxytricha trifallax*

**DOI:** 10.1101/2024.12.31.630814

**Authors:** Danylo J. Villano, Manasa Prahlad, Ankush Singhal, Karissa Y. Sanbonmatsu, Laura F. Landweber

## Abstract

Genome organization recapitulates function, yet ciliates like *Oxytricha trifallax* possess highly-specialized germline genomes, which are largely transcriptionally silent. During post-zygotic development, *Oxytricha*’s germline undergoes large-scale genome editing, rearranging precursor genome elements into a transcriptionally-active genome with thousands of gene-sized nanochromosomes. Transgenerationally-inherited RNAs, derived from the parental somatic genome, program the retention and reordering of germline fragments. Retained and eliminated DNA must be distinguished and processed separately, but the role of chromatin organization in this process is unknown. We developed tools for studying *Oxytricha* nuclei and apply them to map the 3D organization of precursor and developmental states using Hi-C. We find that the precursor conformation primes the germline for development, while a massive spatial reorganization during development differentiates retained from eliminated regions before DNA rearrangement. Further experiments suggest a role for RNA-DNA interactions and chromatin remodeling in this process, implying a critical role for 3D architecture in programmed genome rearrangement.

## Introduction

Genome function depends heavily on chromatin organization. 3D structural features often recapitulate transcriptional or chromatin state and can follow highly dynamic patterns during the cell cycle.^1–6^ Many features persist not just from one generation to the next but also across evolutionary time scales.^7–9^ The nuclear landscape surrounding DNA can be as important to proper genome function as the DNA sequence itself.

A hierarchy of 3D chromatin features has been identified, but the relative importance and universality of these organizing principles are unclear.^10^ Eukaryotic 3D genome structures typically share a dependence on cohesin,^11^ and organize around compartments or compartmental domains based on chromatin state.^9,12,13^ However, varied patterns of chromosome folding have emerged across evolution.^14^ At a smaller scale, topologically-associating domains (TADs)^15^ may enclose conserved genes and persist across cell-type differentiation and speciation^16,17^ and their disruption may lead to cell dysfunction and disease.^3,18^ But the prominence of TADs outside metazoa is an open question,^19–21^ and their size and structure can range considerably, even from cell-to-cell.^22,23^ The factors that shape these structures also vary outside mammalian genomes, in which TAD boundaries often form through the blocking of cohesin-mediated loop extrusion by CCCTC-binding factor (CTCF).^24–26^ Preferential binding of CTCF provides a basis upon which natural selection can act,^27^ but in organisms that lack CTCF,^28–31^ or in those like *Drosophila* where it plays a less central role, a wide range of chromatin-associated proteins mediate structural featues.^32,33^ Compartmental partitioning correlates with transcriptional state but remains even in the absence of transcription;^34^ compartmental domains may form via phase separation, derived from biophysical properties of chromatin.^35^ These previous studies suggest a wide, diverse adaptability in the relationship between genome structure and genome function.

The genome dynamics of the ciliate *Oxytricha trifallax* create unusual demands on genome architecture. *Oxytricha* is a binucleate, unicellular protist with genome functions split between two distinct types of nuclei.^36,37^ All gene expression during vegetative (asexual) growth derives from a somatic genome in a large macronucleus (MAC), comprising over 18,000 highly polyploid, kilobase-scale “nanochromosomes,” most bearing a single gene, two telomeres, and very little non-coding DNA.^38,39^ During sexual reproduction, however, no somatic DNA is inherited from parent to daughter cell. Instead, *Oxytricha* possesses a diploid, transcriptionally-silent germline genome with megabase-scale chromosomes in a micronucleus (MIC), which, at the onset of its sexual lifecycle, undergoes meiosis before fusing with another haploid MIC of a conjugating cell.^40^ A copy of the resulting zygotic MIC develops into a new MAC over 72 hours, while the parental MAC degrades.

Though other transcriptionally silent genomes do demonstrate higher-order, developmentally dynamic 3D patterns,^41^ the *Oxytricha* germline genome must act as the substrate for an intricate program of developmental genome rearrangement that discards roughly 90% of DNA, retaining and joining a set of *macronuclear-destined sequences* (MDSs) that together construct a new somatic genome.^40^ MDSs that are syntenic in the MAC are interrupted by *internally eliminated sequences* (IESs) in the germline, and frequently out of order (“scrambled”) or tens of kilobases apart. During development, hundreds of thousands of MDSs must be recognized, marked for retention, protected from degradation, then recombined in the correct order before amplification to high copy number. Rearrangement occurs between short direct repeats at MDS ends and is guided by two classes non-coding RNA (ncRNA) — 27nt piRNAs and long “template RNAs,” respectively.^42–44^ These ncRNAs derive from the parental MAC and provide transgenerational memory of which sequences to retain and the order in which to link them.

How 3D organization of chromatin facilitates rearrangement of the *Oxytricha* germline is unknown. It is as critical that some regions be degraded as that adjacent sequences be protected, and that syntenic MDSs be in close physical proximity. The complex linear arrangement of MDSs in the germline suggests a key role for chromatin organization in *Oxytricha* genome dynamics, and that understanding 3D architecture may illuminate the process of rearrangement. Studies in other ciliates suggest that histone modifications and chromatin state mediate the distinction between retained and eliminated sequences,^45,46^ and in a distantly-related ciliate with modest levels of rearrangement, *Tetrahymena thermophila*, the boundaries of retained germline sequences form TAD-like structures.^47^ Only a minority of *Tetrahymena* sequences are eliminated, germline MDSs are in closer proximity and rarely disordered, and *Tetrahymena* has far fewer germline or somatic chromosomes than *Oxytricha*.^48^ The role of germline chromatin organization in *Oxytricha*’s complex rearrangement has been unknown, in either the initial precursor state or the developing nuclei (called “anlagen”).

Here we present the higher-order 3D genome structure in the germline of *Oxytricha trifallax* in both its precursor and developmental states. We scaffold the previously fragmented germline genome assembly using 3D genomics and long-read sequencing. Through Hi-C, we identify TAD-like structures in the precursor germline that suggest the genome is poised for rearrangement even prior to development. We then present the wholesale reorganization of the genome at the onset of sexual development, which appears to facilitate rearrangement through spatial separation of eliminated and retained sequences. Additional experiments demonstrate that retained sequences increase in both chromatin accessibility and RNA-DNA interactions during development. These results inform a model in which dynamic chromatin organization plays a critical role in mediating genome rearrangement.

## Results

### Separation of *Oxytricha* nuclei via FACS based on size and DNA content

In order to study the conformation of germline chromatin in the absence of MAC contamination, we first developed approaches to purify different types of *Oxytricha* nuclei. Previous methods for enriching *Oxytricha* germline DNA do not preserve nuclear integrity or provide sufficient purity.^49^ Fluorescence-activated cell sorting (FACS) has recently been used to isolate nuclei of other ciliate species based on nuclear proteins.^50,51^ However, the size and DNA content of *Oxytricha* nuclei differ so vastly that sorting based on DAPI staining permits scalable purification of intact *Oxytricha* nuclei. Suspensions of formaldehyde-fixed asexually growing *Oxytricha* nuclei of strain JRB310 were sorted by FACS on the basis of particle size and DAPI signal. Clusters of putative large, DNA-rich MACs and small, comparatively DNA-poor MICs are readily distinguishable (Figures 1A and S1A). These populations were sorted, and their DNA assessed via qPCR for somatic-specific and germline-specific sequences, which confirmed the identity of each population and demonstrated the lack of MAC contamination in the MIC sample (Figure 1B). The expected difference in molecular weight of DNA extracted from each population was also confirmed (Figures S1B and S1C).

**Figure 1.**
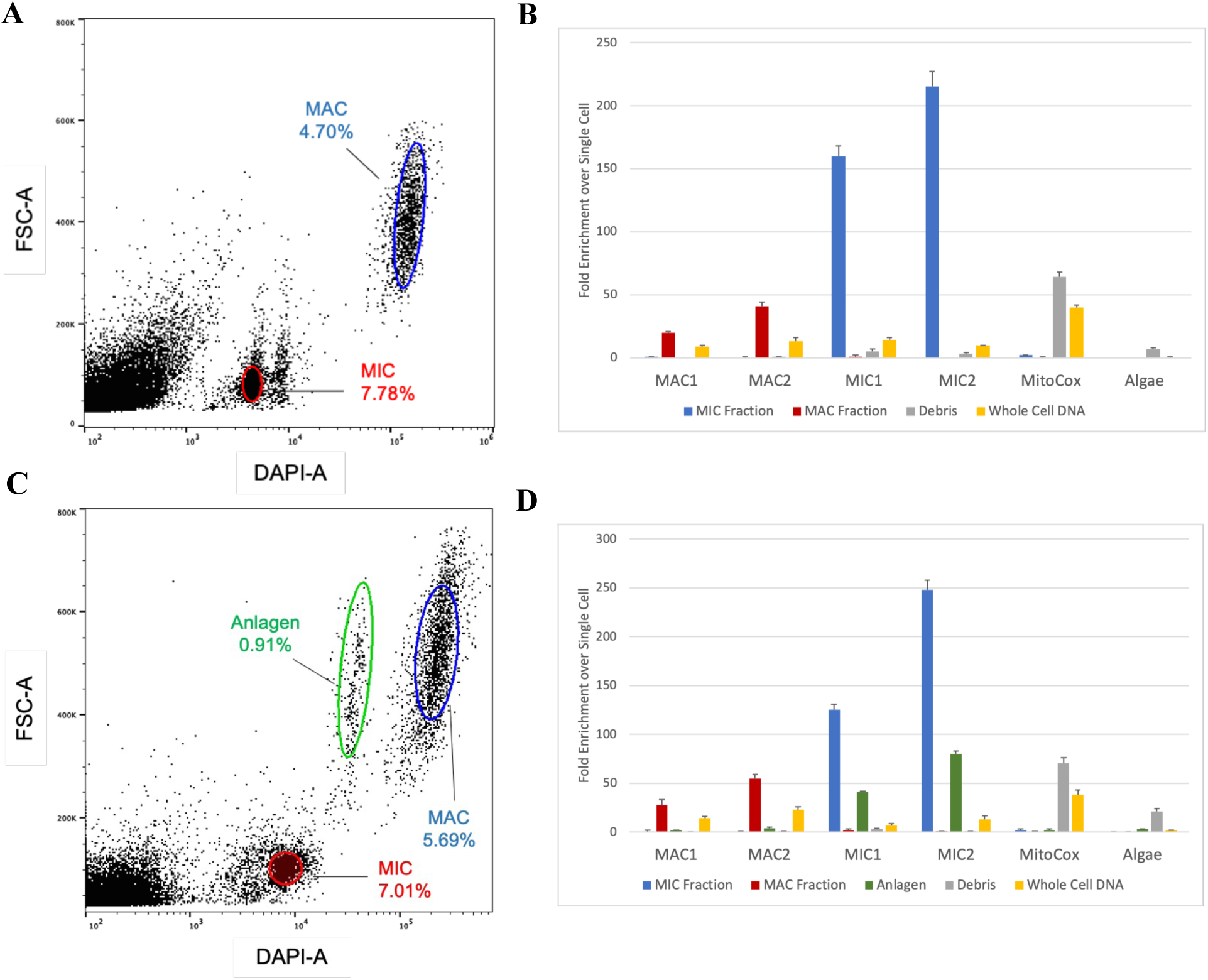
Isolation of *Oxytricha* nuclei via FACS. (A) Selection of vegetative JRB310 nuclei based on DAPI and forward scatter (“FSC”) signal. (B) qPCR of sorted vegetative populations. “MAC1” and “MAC2” target MAC-specific loci that span scrambled MDS-MDS junctions. “MIC1” and “MIC2” target high copy number MIC-specific loci within transposable elements. “MitoCox” targets the mitochrondial COX1 locus. “Algae” targets *Chlamydomonas reinhardtii*-specific sequence. “Debris” sample derives from dense, undefined cloud of particles to the left of MIC population in (A). (C) Selection of developmental nuclei at 24h post-mixing of JRB310 and JRB510 cells. (D) qPCR of sorted developmental populations, as in (B).

Suspensions of post-zygotic, developing *Oxytricha* nuclei were prepared and sorted in the same manner. Nuclei were harvested at 24 hours post-mixing of mating-compatible cells (24h), a timepoint early in development when germline DNA in the anlagen has been polytenized to approximately 32X^52^ but most DNA rearrangements have not yet begun,^53^ and anlagen have expanded to nearly the size of the old MAC.^42^ Meanwhile, degradation of the old MAC has not yet begun, leading to the observation of all three types of nuclei, using the same size and DAPI signal criteria as before (Figures 1C and S1D). These developmental nuclei were tested via qPCR using the same MIC-and MAC-specific sequences as before, confirming the presence of germline-specific DNA in the population of anlagen (Figure 1D).

### 3D and long-read scaffolding improves contiguity of the reference germline genome

Previous studies estimated 75-120 megabase-scale chromosomes in the *Oxytricha* germline genome.^52,54^ However, the published JRB310 germline reference genome, based on sequencing of size-selected DNA from MICs enriched via sucrose gradient, has more than 25,000 contigs, all less than 500 kb, covering a total of 496 Mb of sequence.^40^ The previous inability to generate larger, chromosome-sized contigs derived from both the impurity of sucrose gradient-enriched MIC DNA^49^ and the repetitiveness of the germline, which consists of over 30% repetitive sequences alongside extensive MDS paralogy.^40,55^ In order to evaluate 3D germline features, a reference of much higher contiguity than was previously available was needed.

We took a combined approach to scaffold the *Oxytricha* germline into larger contigs, using 3D genomics and long-read sequencing. Approximately 1µg genomic DNA was extracted from two biological replicates of FACS-sorted vegetative MICs, which were then sequenced via Oxford Nanopore (Figure S2).^56^ In parallel, *in situ* Hi-C libraries were built (Figure S3) from two biological replicates of over 1,000,000 FACS-sorted vegetative MICs,^57^ which were then sequenced via Illumina, yielding >500M paired-end reads. When Hi-C reads were mapped to the reference germline genome using Juicer,^58^ the highly fragmented state of the assembly was clear, as numerous small contigs interacted with one another at very high frequency, suggesting they were in fact located on the same chromosome (Figure 2A). To improve the assembly, we first used LRScaf to bridge original contigs using germline Nanopore data (Figure S4A).^59^ Next, Hi-C reads were used to scaffold the product of LRScaf using 3D-DNA, based on the principle that intrachromosomal contacts are far more likely than interchromosomal contacts.^60^ Output from 3D-DNA was visually inspected using Juicebox^61^ (Figure S4B), and obvious errors produced by Hi-C-based scaffolding were manually corrected in that program. 3D-DNA leaves scars of ambiguous bases between scaffolded contigs, so another round of LRScaf was performed for correction.

**Figure 2.**
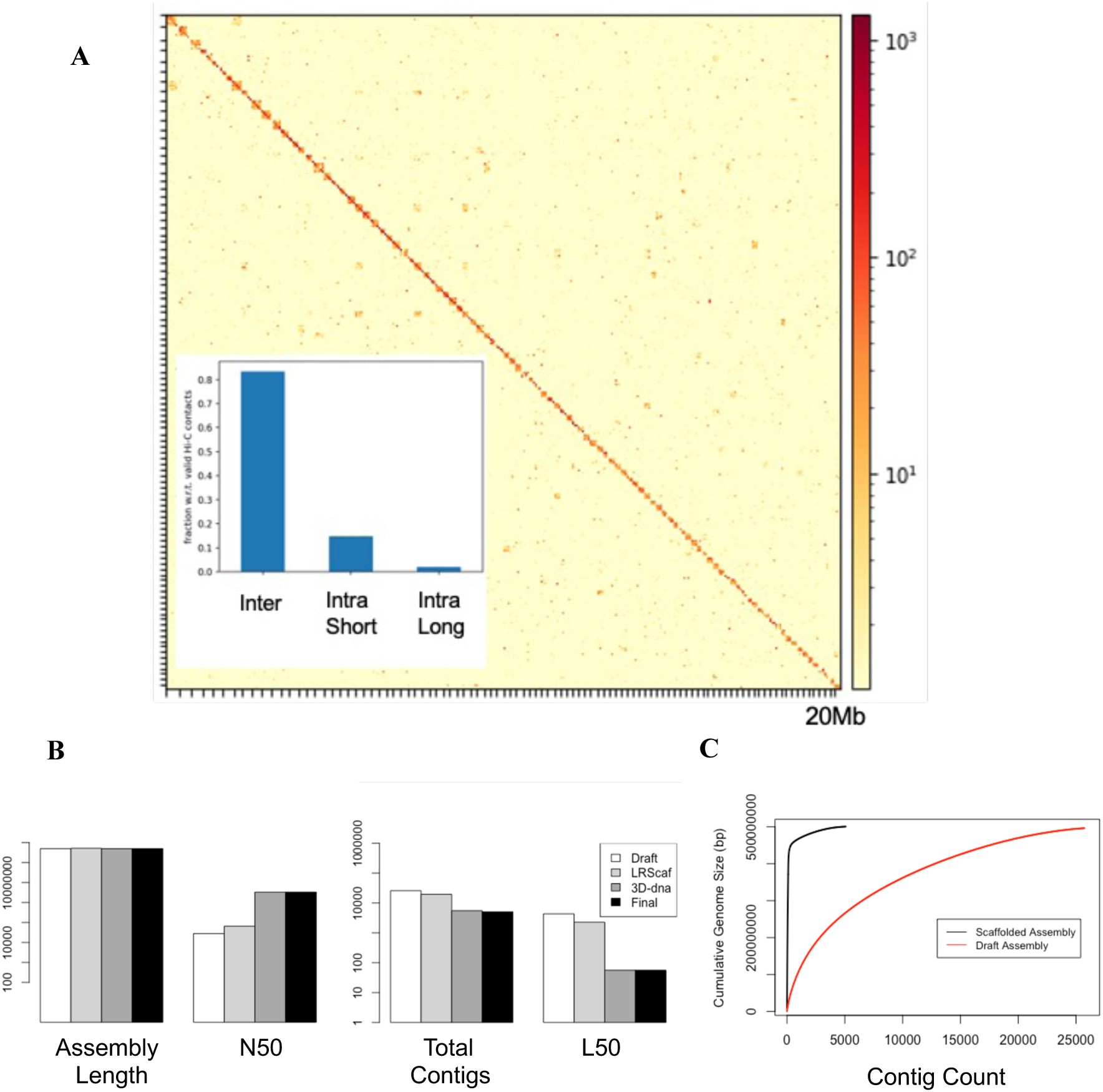
Hi-C and long-read data produce a highly contiguous scaffolded genome assembly. (A) Hi-C map of vegetative germline Hi-C data (log1p) mapped to the original assembly’s 100 longest contigs, totaling ∼4% of the genome and <0.5% of contigs. Hash marks indicate contig boundaries. High interaction frequency between small contigs indicates likely synteny. Inset shows frequency of different interaction types: Interchromosomal, Intrachromosomal Short (≤ 20kb), and Intrachromosomal Long (>20kb). (B) Assembly statistics after different stages of scaffolding. “Draft” indicates original germline reference genome.^40^ (C) Cumulative distribution of contig sizes in original and scaffolded assemblies.

The final scaffolded assembly has substantially improved contiguity and 12 Mb additional sequence (Figure 2B). The initial 25,720 contigs is reduced nearly 5-fold. Despite the total contig count remaining over 5,000, most of these are very small and unable to be processed by 3D-DNA. 81% of the total genome is present on 123 contigs larger than 1 Mb, while 90% of the genome is contained within 358 contigs larger than 50 kb (Figures 2C and S4C). Two megabase-scale scaffolds contain telomeric repeats at both ends (Table 1). Aligning known MDSs from the somatic genome to the scaffolded germline genome via BLAST^62^ produced a similar mappability profile as when aligned to the original assembly (Figure S5A). Upon remapping vegetative germline Hi-C data to the scaffolded assembly, the number of intercontig interactions was reduced more than 40%, reflecting that most were actually long-range (>20 kb) intrachromosomal interactions. There remains an excess of interchromosomal contacts genome-wide, consistent with the current level of scaffolding compared to the estimated ∼75 chromosomes in the MIC. If we consider only contacts with at least one read pair mapping to the thirteen scaffolds >5 Mb, then interchromosomal contacts are reduced more than 69% relative to the original fragmented assembly (Figure S5B). This more complete understanding of the 1D architecture of the *Oxytricha* germline now permits analysis of its 3D architecture.

**Table 1.**
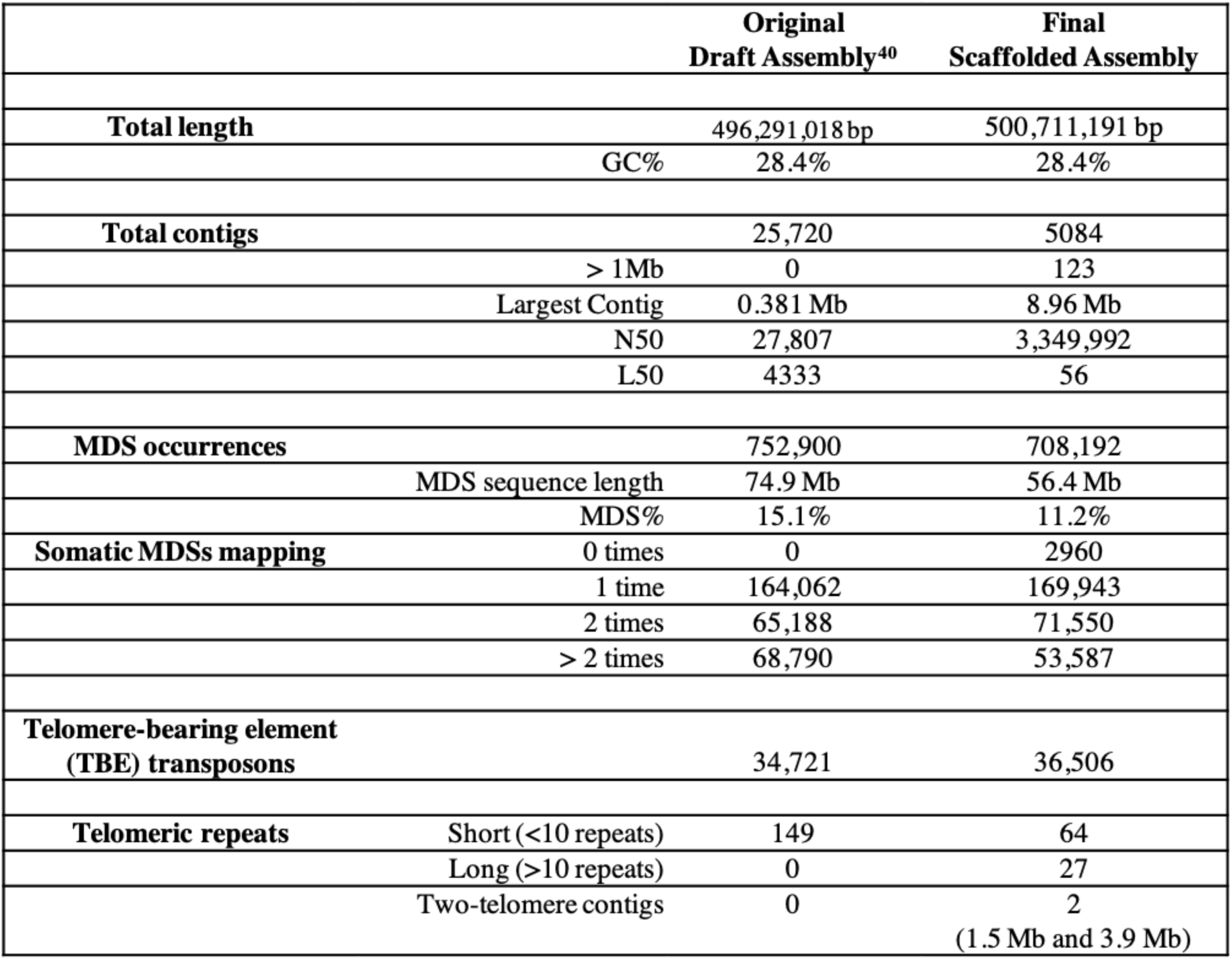
Germline micronuclear genome scaffolding statistics.

### TAD-like structures in the precursor germline cluster retained sequences

In order to learn how MDSs are organized in vegetative micronuclei, germline Hi-C data mapping to 123 scaffolds longer than 1 Mb (covering 81% of total germline sequence) were extracted from the contact matrix and analyzed with HiCExplorer (see Methods). After KR matrix balancing,^63^ from a genome-wide view there was little interchromosomal structure (Figures 3A and S6A), with scaffolds resembling classical chromosome territories^64^ and low compartmentalization (Figure S6B). Long-range loops could not be reliably detected in this largely heterochromatic genome,^36^ due in part to the resolution of the data. Calling TADs at resolutions of 5, 10, and 100 kb, however, revealed the presence of dozens of TAD-like structures (Figure 3B).

**Figure 3.**
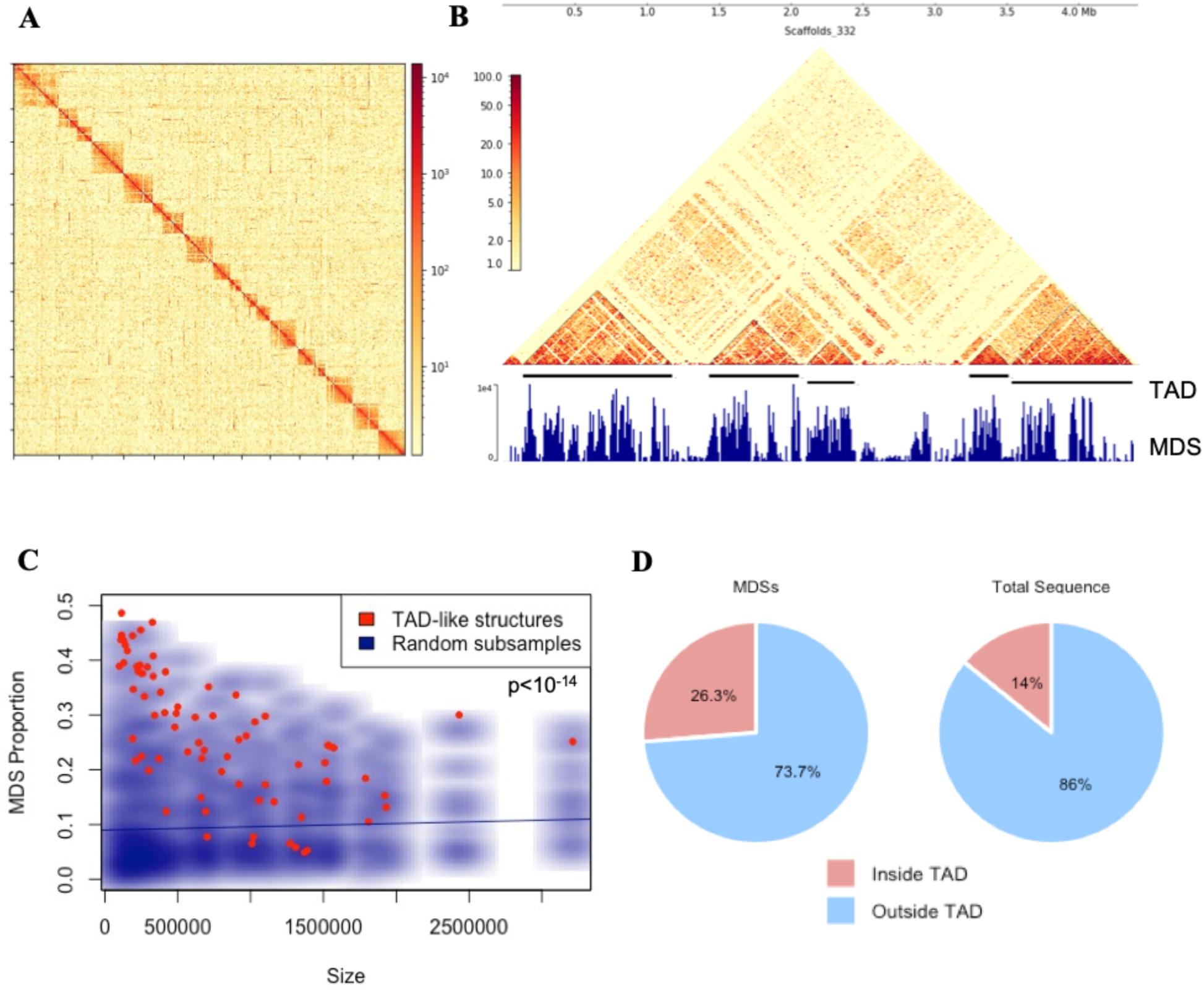
TAD-like structures form around dense tracts of retained sequences. (A) Germline Hi-C contacts (log1p) from 13 scaffolds larger than 5 Mb, totaling 78 Mb. (B) Germline Hi-C contacts (log1p) from representative scaffold bearing several TAD-like structures. “MDS” track shows total MDS sequence within 10 kb bins. (C) Distribution of TAD-like structure lengths and MDS proportion compared to smoothed scatter plot of 25 iterations of randomly subsampled regions of the genome with the same size distribution as the TAD-like structures. Line is linear model of the random genomic subsamples. *p*-value calculated by Student’s *t*-test on MDS proportion of TADs vs. average of subsample iterations. (D) Percent of total MDS sequence and genomic sequence contained within TAD-like structures.

Ranging in size from 100 kb to 3.2 Mb, these domains were found to be highly enriched in MDSs (Figure 3C; p<10^-14^, Student’s *t*-test), which tend to occur in dense tracts in the germline (Figure S7), such that terminal MDSs of different nanochromosomes often overlap.^40^ The TAD-like structures have a median MDS content of 26%, compared to 11% for the germline as a whole (Figure S6C). Only 7/74 TAD-like structures had less than the genome-wide average MDS content. After splitting the entire germline genome into 100 kb regions, 46% of the ≥ 90^th^- percentile most MDS-rich bins at least partially overlapped with the TAD-like structures, including 20 of the 30 richest bins (Figure S6D). In total, over 25% of MDSs were contained within these subchromosomal 3D features (Figure 3D).

TADs are preferentially self-associating regions of chromatin; sequences within TADs are more likely to interact with one another than expected based purely on their 1D distance. In transcriptionally active genomes, TADs permit genes to make contact with distant enhancers.^65^ The *Oxytricha* germline has no need of enhancers, but analogously, its TAD-like structures similarly cluster sequences that benefit from proximity. Co-localization of MDSs reduces the nuclear space in which rearrangement machinery needs to find its substrates, and may facilitate specific interactions between distant MDSs that ultimately will be linked. The 3D co-localization of *Oxytricha* MDSs approximates the linear clustering of MDSs throughout the genome, and suggests that germline chromatin organization anticipates the spatial needs of genome rearrangement, even in the absence of development.

### Chromatin dramatically reorganizes during development, distinguishing retained and eliminated regions

We then investigated how 3D conformation changes with the onset of development to permit wholesale genome rearrangement. Nuclear development requires approximately 72 hours to complete. At 24h, the genome is on the cusp of rearrangement. ncRNAs that program genome rearrangement are abundant in the anlagen,^42,66^ and germline DNA has been polytenized to ∼32X,^52^ but massive DNA editing has not yet occurred.^37^ Two biological replicate Hi-C libraries (Figure S8) were prepared from > 250,000 FACS-purified 24h anlagen. Illumina sequencing produced 570 million paired-end reads, which were mapped to the scaffolded reference genome with Juicer,^57^ then corrected and analyzed with HiCExplorer.^31,67,68^

24 hour Hi-C contacts across all scaffolds ≥ 1 Mb revealed a striking shift in 3D genome architecture (Figure S9A and S9B). Among the largest scaffolds, *inter*chromosomal contact frequency tripled compared to the precursor, vegetative state, along with an enormous (>5-fold) reduction in the frequency of long-range *intra*chromosomal contacts > 20kb (Figures S9C and S9D). With the loss of precursor chromosome territory-like organization, the structure of vegetative TAD-like structures was also largely absent, and fewer than 11% could be called at 24h (Figure S10). We instead detected the emergence of a dramatic 3D architectural bifurcation of the genome. Whereas there were no apparent compartments in the vegetative germline, the signature checkerboard pattern reflecting compartmentalization^69^ arises in the 24h anlagen (Figure 4A), confirmed by compartmentalization analysis (Figure 4B).^70^ The 3D profile of the two compartments is starkly different. While interchromosomal interactions are enriched genome-wide at 24h, they are 44% more frequent in compartment A vs. compartment B bins (based on the sign of the first eigenvector) (Figure 4C). Inter-and intrachromosomal interactions are roughly equal in compartment B, but in compartment A, which is highly enriched in MDSs, nearly 90% of contacts are *inter*chromosomal. Though the genome is roughly equally divided between each compartment (51% vs. 49%), retained segments are more than twice as likely to be located in compartment A than compartment B (Figure 4D). Unlike in the vegetative germline, MDS content correlates well with PCA1, and over 80% of the 30 most MDS-enriched 100 kb bins are found in compartment A (Figures S11A and S11B). DNA fluorescence *in situ* hybridization (Figure S12)^71,72^ and multi-chromosomal 3D reconstructed models (using a modified version of the 4DHiC method,^73^ allowing for multiple chromosome interactions^74^) of the experimental Hi-C data (Figure S13)^75–77^ corroborate nonuniform subnuclear localization of developmental compartments and increased interchromosomal mixing during development relative to the precursor vegetative state.

**Figure 4.**
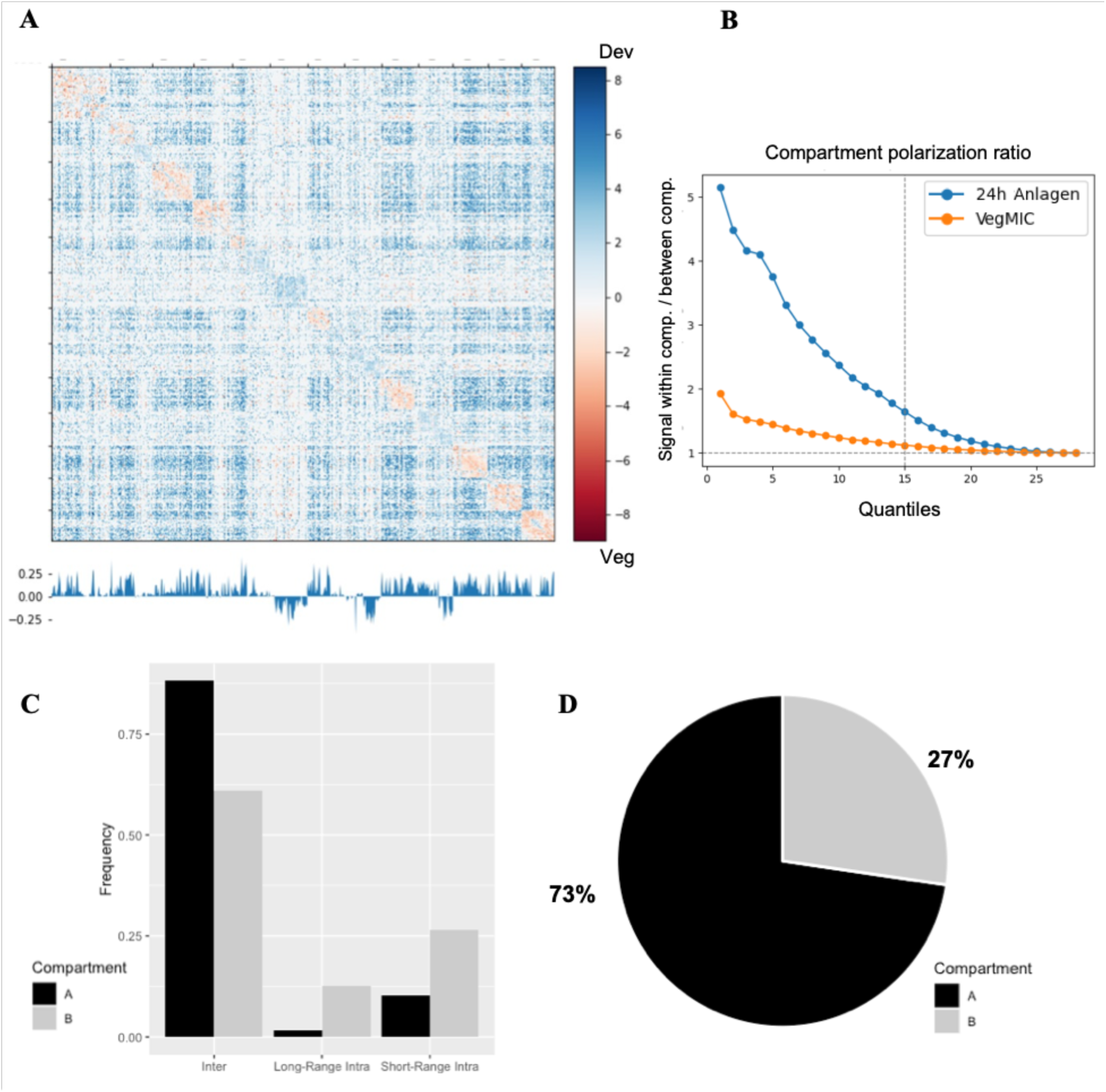
The developing genome compartmentalizes to separate retained and eliminated regions. (A) 24h anlagen Hi-C data across scaffolds ≥ 5 Mb. Plotted values are log_2_ ratio over vegetative germline (input matrices KR-corrected and normalized by coverage). Track below is PC1 value indicating compartment. (B) Compartmentalization signal of the 24h developmental genome relative to vegetative germline. 24h PC1 values used for division into quantiles. (C) Frequency of different interaction types within compartment A or B bins at 24h. (D) Percentage of MDS sequence in compartment A or B at 24h.

We then investigated whether this radically altered 3D architecture persisted later into genome rearrangement. At 48h, massive DNA editing is underway and rearrangement intermediates are abundant, as somatic sequences are rebuilt before the final stage of DNA amplification. Hi-C was performed on ∼80,000 sorted 48h anlagen. When sequences were aligned and analyzed as before, despite lower coverage than the previous libraries (Table S1), the 48h anlagen present marked changes (Figures S14A and S14B). A checkerboard pattern can still be discerned when compared to the vegetative germline (Figure S14C), but weaker than at 24h (Figure S14D). The frequency of interchromosomal contacts (Figure S9C), the compartmentalization signal (Figures S15A and S15B) and the correlation of PCA1 with MDS content (Figure S11C) were all intermediate between the precursor germline and the 24h anlagen. Although the 48h late developmental genome shares with the 24h early developmental genome a dearth of long-range *intra*chromosomal contacts relative to the germline, and although the developmental samples correlate well (Figure S15C), the 48h developmental genome shows a striking increase in short range (< 20 kb) contact frequency, twice that of the precursor germline (Figure S9C), largely at the expense of interchromosomal contacts (Figure S15D). The ongoing deletion of germline-limited regions at 48h complicates this analysis, but this late-stage focus on short-range contacts agrees with the timeline of rearrangement. Somatically syntenic sequences tend to cluster in the germline,^40^ and by 48h the ncRNA-guided joining of consecutive MDSs is underway. Earlier, at 24h, the reorganization of the germline genome into compartments drives MDSs non-specifically across the genome into a shared nuclear space, with MDSs marked for retention and protected from degradation.^42^ As genome rearrangement progresses during development, the 3D architecture transitions from bulk reorganization to promoting smaller-scale interactions.

### Retained and eliminated sequences are distinguished by chromatin state

The widespread shift in germline 3D genome architecture during development must ultimately distinguish retained vs. eliminated sequences, allowing the former to interact with each other and insulating the latter. Such chromatin distinctions are a hallmark of differences between compartments in metazoan genomes,^26^ and have been observed during development in other ciliates.^45,78^ Since disparities in chromatin accessibility are a primary functional difference between chromatin states,^79^ we measured chromatin accessibility in the *Oxytricha* germline by performing ATAC-seq^80^ on sorted nuclei from vegetative and 24h developmental cells (see Methods).

Focusing on features of intermediate size, we observed early in development that retained regions, i.e. MDSs, were far more accessible than deleted regions (IESs; Figure 5A; p < 10^-10^, Student’s *t*-test). This relationship was present in the precursor germline as well, as IESs were nearly absent in the germline ATAC-seq data, but MDSs are much more accessible during development. ATAC-seq corroborated the developmental bifurcation of the genome, with 100 kb bins in compartment A averaging nearly twice the accessibility of bins in compartment B at 24h (Figure 5B; p<10^-13^, Student’s *t*-test). These regions had little difference in the precursor MIC. Peak calling in ATAC-seq coverage displays MDS enrichment in both vegetative and developmental samples, though moreso in the latter (Figure 5C). Curiously, telomere-bearing elements (TBEs) — semi-domesticated transposable elements that are transiently expressed from the developing anlagen at 24h before self-elimination^81,82^ and represent approximately 15% of the MIC genome^40^ — are also highly enriched in anlagen peaks and depleted in the vegetative data. Averaging the coverage over thousands of germline features reveals the precision of these differences in accessibility. In both states, accessibility increases sharply at MDS boundaries (Figure 5D); conversely, accessibility drops precisely within IESs (Figure 5E). Likewise, TBE coverage spikes at either end of the transposon during development (Figure 5F), which may reflect its active transcription.^82^ These results suggest a clear contrast between the chromatin state of retained vs. eliminated sequences in the *Oxytricha* germline, accompanying their 3D separation.

**Figure 5.**
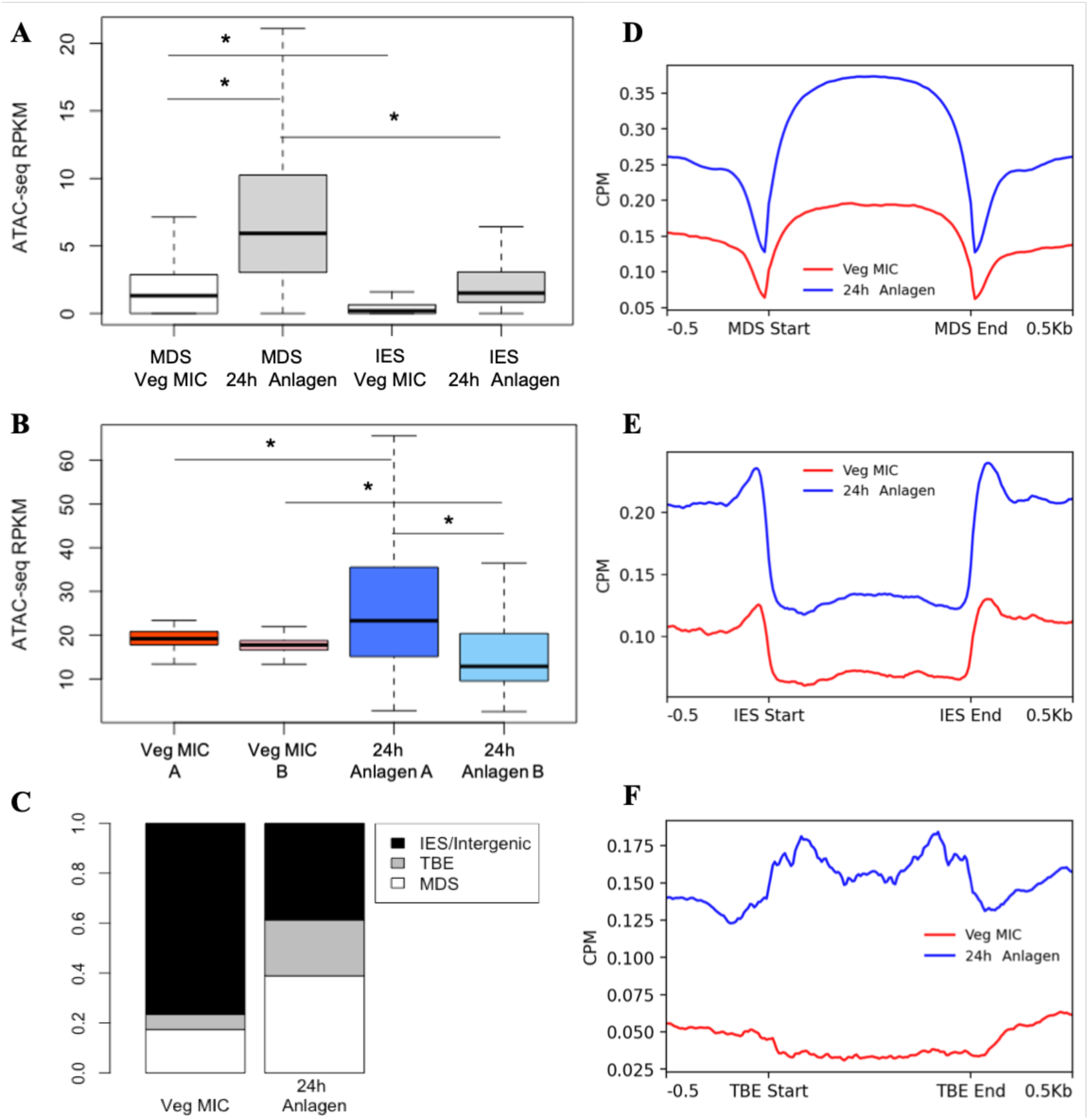
Retained sequences are more accessible at 24h than eliminated sequences. (A) ATAC-seq coverage over MDSs and IESs in vegetative germline and 24h anlagen. Germline features were restricted to those of similar size, between 500bp-1,000 bp (> 10,000 features for each type). * indicates *p*-value <10^-10^, Student’s *t*-test. (B) Vegetative and developmental ATAC-seq coverage within 100 kb regions identified as compartment A or B at 24h. * indicates *p*-value <10^-13^, Student’s *t*-test. (C) Proportion of vegetative and 24h ATAC-seq peaks mapping within MDSs, TBEs, or other eliminated sequences. (D) ATAC-seq coverage profile of size-selected MDSs in each ATAC-seq dataset. (E) ATAC-seq coverage profile of size-selected IESs in each ATAC-seq dataset. (F) ATAC-seq coverage profile of size-selected TBE transposons (3-4 kb) in each ATAC-seq dataset.

### RNA mediates long-range DNA interactions during development

As in other ciliates, sequence retention and elimination in *Oxytricha* development is regulated by transgenerationally-inherited ncRNA.^83^ piRNAs and template RNAs, both of which derive from genome-wide transcription of the parent’s somatic genome, peak in expression prior to the 24h timepoint at which we observe compartmentalization.^42,44^ The specific mechanism by which these ncRNAs lead to retention is unknown, but their precise targeting of retained sequences makes them strong candidates to contribute to the developmental spatial separation between MDSs and eliminated regions. Several recent studies have demonstrated the influence of ncRNA on 3D genome architecture, through either binding of nascent transcripts or RNA-DNA interactions.^84–86^ In light of the massive processing of all MDSs and the lack of widespread transcription from the anlagen,^66^ we conjectured that ncRNA may be mediating germline development through direct RNA-DNA interactions.

In order to test whether RNA may be directly binding MDSs during development and thereby supporting chromatin reorganization, we measured RNA-DNA hybrid abundance using the S9.6 antibody (Figure 6A).^87^ We observed increased levels of RNA-DNA hybrid formation during *Oxytricha* development, relative to the precursor, vegetative germline, peaking at 24h and persisting through the end of genome rearrangement. To map the locations of these hybrids, DNA-RNA immunoprecipitation sequencing (DRIP-seq)^88^ was performed on whole cell DNA from precursor, vegetative cells and developing *Oxytricha*. The resulting reads were mapped to the somatic genome, which provides the source of developmental ncRNA.^42,43^ Before development, hybrids in asexually dividing cells were found to be enriched at both the 5’ and 3’ ends of single-gene nanochromosomes (Figure 6B). Because transcription start sites (TSS) and termination sites (TTS) are very close to nanochromosome ends,^54^ these hybrids are likely transcription-associated, as transcriptional R-loops tend to form at the TSS and TTS.^89^ However, DRIP-seq at the 24h timepoint shows elevated coverage across all MDSs, regardless of distance from nanochromosome ends (Figure 6B). This ubiquitous hybridization of RNA across retained MDS sequences during development suggests direct hybridization of MAC-derived ncRNA to the developing, anlagen DNA.

**Figure 6.**
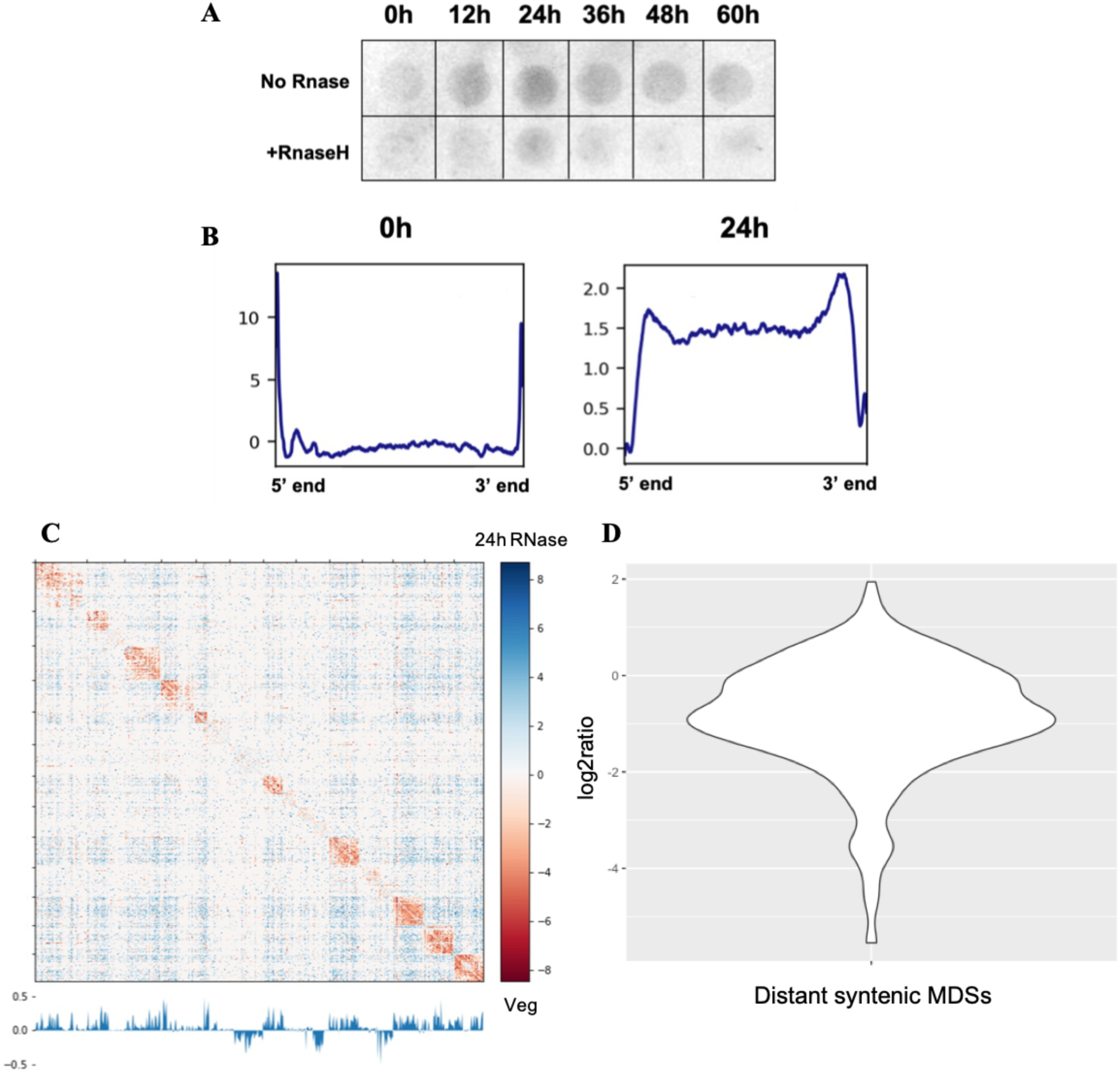
RNA-DNA interactions influence developmental 3D organization. (A) Dot blot of RNA-DNA hybrids across developmental timecourse visualized with S9.6 antibody, including RNase H control. Remaining signal in RNase H control is most likely double-stranded RNA, which has some antibody affinity^100^ and are ncRNA precursors in developmental.^43^ Membrane loaded with 130ng genomic DNA per sample. (B) DRIP-seq coverage across 5,000 nanochromosomes (all 1.5-3 kb, bearing single genes) in vegetative and developmental states. (C) Log_2_ ratios of RNase-treated 24h anlagen Hi-C vs. vegetative germline across scaffolds > 5 Mb. Lower track is PCA1 value of RNase-treated 24h anlagen in 100 kb bins. (D) Violin plot of log_2_ ratios of contact frequency between 10 kb bins bearing distant, syntenic MDSs, RNase-treated 24h anlagen/24h control.

To test whether RNA provides scaffolds to assist the developmental compartmentalization of the *Oxytricha* germline genome, we treated 24h developing cells with a robust RNase treatment to digest the RNA strand in RNA-DNA hybrids, as well as all single-and double-stranded RNA, prior to fixation and FACS, alongside control, permeabilized cells that were not RNase treated. Hi-C on RNase-treated, sorted anlagen still displayed a qualitatively similar 3D checkerboard pattern relative to the control (Figure 6C), indicating that compartmentalization was not strongly affected by the treatment (Figures S16A and S16B). This suggests that by 24h, RNA is no longer critical for the overall maintenance of this chromatin superstructure. Though interchromosomal interactions are still elevated after RNase treatment, relative to the precursor germline, long-range intrachromosomal interactions were reduced more than twofold relative to the control (Figure S16C). Since template RNAs derive from somatic nanochromosomes, and most MDSs that form the same nanochromosome are syntenic in the germline, we hypothesized that RNA could be mediating specific interactions between precursor MDSs that contribute to the same MAC chromosome. Binning the Hi-C data at 10 kb, we identified the 400 most distant pairs of MDSs that are located on the same contig in both the somatic and germline genomes, since enrichment of interactions between these genomic bins are most easily measurable. Comparing RNase-treated anlagen to the control, we found an average of almost twofold reduction in the frequency of these distant intrachromosomal interactions, and over 75% of cases decreased with RNase treatment (Figure 6D). This result, along with the increased level of RNA-DNA hybrids, supports a model in which RNA may directly scaffold rearranging DNA^44^ to facilitate the colocalization of MDSs that recombine to form a mature nanochromosome during development, exploiting the broader context of 3D chromatin architecture.

## Discussion

*Oxytricha trifallax* offers a direct yet unconventional relationship between genome structure and function. We show that dramatic chromatin reorganization precedes massive genome rearrangement. The dynamic architecture of its genome, with transcriptionally-silent and often scrambled, precursor gene segments representing the functional units of DNA, foreshadows its extensive remodeling. MDS-rich 3D domains in the precursor germline micronucleus poise the genome for development, and then extensive compartmentalization spatially separates predominantly retained from eliminated regions early in development, followed by a coalescence of short-range interactions late in development, when new macronuclear chromosomes form.

In the distantly-related ciliate *Tetrahymena,* with considerably less genome rearrangement, the boundaries of germline TAD-like structures coincide with chromosome breakage sites during development,^47^ and in mammalian systems, the boundaries of a majority of TADs are demarcated by CTCF-binding sites.^90–92^ *Drosophila* has several DNA-binding insulator proteins that together fill a similar role,^31^ but studies in representative plants, fungi, and protists with higher order chromatin structure have yet to identify functionally similar architectural proteins.^30,93^ *Oxytricha*, with orders of magnitude more breakage sites than *Tetrahymena*, also lacks a CTCF homolog; however, other master regulators of 3D genome architecture may play a prominent role in its genome dynamics.^94^

Compartmentalization of *Oxytricha*’s germline genome during development appears to self-organize around the location of retained sequences, as MDSs are driven into shared nuclear spaces that exclude MDS-depleted regions. This reduction in complexity would encourage productive searching for and joining of consecutive MDSs. It is also unknown whether interactions occur between multiple or single copies of polytenized germline chromosomes. The ATAC-seq data suggest that compartmentalization could depend on homotypic chromatin state interactions. Histone or DNA modifications can be especially dynamic during meiosis and development^95^ and limited histone data are available for the ciliate *Stylonychia*,^45,96^ but little is known about their role in *Oxytricha* development.

The role of ncRNA in *Oxytricha* genome reorganization is particularly intriguing, as two classes of ncRNA provide a code for sequence retention.^42–44^ How ncRNA specifically transmits the information it carries to program genome editing is a major question in *Oxytricha* biology. Like bridge RNAs,^97^ *Oxytricha*’s long template RNAs may provide a scaffolding agent for the rearranging genome, directly mediating thousands of long-range DNA-DNA interactions, while piRNAs may act indirectly to guide changes in MDS chromatin state. *Oxytricha* stands out among the main ciliate model systems in that its largest class of small RNAs mark retained rather than eliminated sequences, although MDSs occupy a smaller fraction of the *Oxytricha* genome, and in each species the most abundant class of small RNAs typically targets the minority class of sequences for DNA retention or deletion.^42^ For example, in *Tetrahymena*, scnRNA targeting of eliminated sequences leads to histone H3 lysine-9 methylation,^46^ which can cause heterochromatin formation in an RNA-guided manner.^98^ In *Oxytricha*, piRNA targeting of retained regions could lead to a chromatin mark that prevents heterochromatinization and promotes chromatin accessibility and compartmentalization, ultimately ensuring retention.

Hi-C, itself, has limitations, starting with the depth of DNA sequencing, the complexity of DNA recovered after sorting nuclei, and the inability to uniquely map short reads across a complex germline genome. Future work may improve the yield of FACS-based nuclear sorting, incorporate multimapping reads through probabilistic mapping, or obtain higher resolution through variations like Micro-C.^93^ Currently, we must draw inferences on how MDSs are organized without the ability to analyze their specific pairwise interactions or call loop contacts, instead relying on binning the genome and analyzing the enrichment of MDSs in specific regions. Future work may confirm whether all or merely a large majority of MDSs are driven into dense colocalization early in development.

Plasticity is also a biological feature of *Oxytricha* development, sometimes permitting combinatorial rearrangement of MDSs to produce new genes.^99^ Polytenization of germline DNA may provide more opportunity for successful rearrangements, as well as room for innovation.

Moreover, the compartmental distinction between retained vs. eliminated sequences does not account for the most granular requirements of genome rearrangement. Long stretches of eliminated sequence can be insulated and segregated from retained sequences by compartmentalization early in development, but how are the shortest IESs identified and removed if they are no larger than introns and interspersed between MDSs in compartment A? Multiple waves of DNA elimination might use different molecular mechanisms for depletion of megabase-long stretches of germline-restricted DNA versus precise excision of small IESs.

While a full understanding of *Oxytricha* genome rearrangement is still coming to light, our work demonstrates a tight-knit relationship between genome structure and function, and how adaptable and flexible that relationship can be in the face of complex DNA dynamics.

## Acknowledgements

We thank Sheela George and Ryan Tran for laboratory support. We thank David Dai, Eoghan Harrington, John Beaulaurier, Lynn Ly, and Sissel Juul at Oxford Nanopore Technologies in New York for providing sequencing and advice. We thank Michael Kissner at the Columbia Stem Cell Initiative Flow Cytometry Institutional Core Facility for providing FACS training and advice. We also thank Brian Beliveau, Bill Jack, Stavros Lomvardas, Lorraine Symington, and all current and past Landweber lab members for discussions about genome architecture and *Oxytricha* biology. This study used the Confocal and Specialized Microscopy Shared Resource of the Herbert Irving Comprehensive Cancer Center at Columbia University, funded in part through NIH/NCI Cancer Center Support Grant P30CA013696. This study was supported by NIH R35-GM122555 and NSF 1764366 to LFL. DJV also received support from 5T32GM008798. KS and AS are grateful to generous support from Los Alamos National Laboratory Institutional Computing for allocations on LANL Chicoma, and LANL Venado. KS and AS thank the generous support from NSF (MCB 2337393), DOE LANL LDRD (20210082DR and 20210134ER) and the U.S. Department of Energy, Office of Science, through the Biological and Environmental Research (BER) and the Advanced Scientific Computing Research (ASCR) programs under contract number 89233218CNA000001 to Los Alamos National Laboratory (Triad National Security, LLC).

## Author Contributions

DJV designed, performed, and analyzed most of the experiments. MP contributed to experiments on RNA-DNA hybrids. AS and KS designed and performed coarse-grained modeling of Hi-C data. DJV and LFL conceived the study and wrote the paper with input from other authors.

## Methods

### *Oxytricha* culturing and mating

Vegetative (asexually growing) *Oxytricha trifallax* strains JRB310 and JRB510 were cultured as previously described.^38^ Cells were maintained at approximately 10,000 cells/ml in Pringsheim media (0.11mM Na_2_HPO_4_, 0.08mM MgSO_4_, 0.85mM Ca(NO3)_2_, 0.35mM KCl, pH 7.0), filtered daily through cheesecloth, and fed daily with *Chlamydomonas reinhardtii*. Cells were also fed every other day with *Klebsiella pneumoniae* liquid cultures. Developmental samples were prepared by mating starved JRB310 and JRB510 at a concentration of 5,000 cells/ml, with timepoints measured from the moment parental cells were mixed (0h). Matings were confirmed to have at least 70% efficiency by counting conjugating pairs.

### FACS sample preparation and sorting

Nuclear suspensions for sorting via FACS were collected by filtering cultures through cheesecloth and concentrating on a 10μm Nitex mesh filter before being pelleted by centrifugation for 2 minutes at 200xg. Cells were washed once in Pringsheim media, then collected again by centrifugation. Cells were then fixed in a 1% formaldehyde (v/v) solution in Pringsheim at room temperature with gentle shaking for 10 minutes. Glycine was then added to a final concentration of 125mM to quench fixation, followed by incubation at 4°C for 15 minutes with gentle shaking. Fixed cells were collected by centrifuging for 2 minutes at 300xg. Cells were then lysed by addition of 50ml of lysis buffer (20mM Tris-HCl (pH 6.8), 3% sucrose (w/v), 0.2% Triton X-100 (v/v), 0.01% spermidine-trihydrochloride (w/v)) and incubation on ice for 5 minutes. Nuclei were collected by centrifugation for 5 minutes at 4,000xg, and then washed in 50ml of FACS buffer (1x PBS, 1mM EDTA, 1% BSA) and collected again by centrifuging for 5 minutes at 4,000xg. Nuclei were resuspended in 1ml FACS buffer per million cells in original culture, and DAPI was added to samples at a 1:1,000 dilution. Samples were stored at 4°C until sorted.

FACS was performed on BD FACSAria II (Becton Dickinson) and MA900 (Sony) machines. FSC threshold was set at 5%, and sensor gain settings were typically as follows: FSC: 5, BSC/SSC: 23%, DAPI: 34%. Sensor settings were adjusted from these benchmarks depending on the signal of individual samples. Gating was first performed on FSC-A x FSC-H to exclude doublets and then on FSC-A x SSC-A to enrich nuclei from cellular debris. Populations of nuclei were then selected on the basis of DAPI-A x FSC-A signal. Sorted nuclei for subsequent sequencing experiments were flash frozen in liquid nitrogen and stored at-80°C.

To confirm the identity of sorted nuclei, DNA was isolated through phenol-chloroform extraction. The difference in the molecular weight of DNA extracted from putative MICs and MACs was measured by BioAnalyzer (Agilent Technologies) and a 1.5% agarose gel. The presence of MAC-and MIC-specific sequences, as well as mitochondrial and algae-derived sequences, in DNA derived from sorted nuclei was then measured via qPCR with Power SYBR Green Master Mix (Thermo Fisher Scientific) on a Bio-Rad CFX384 machine using the following primers:

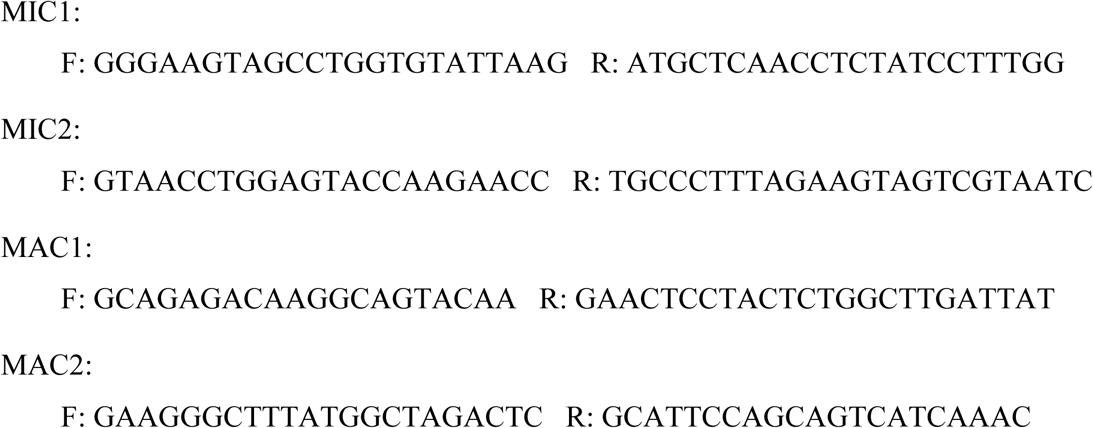

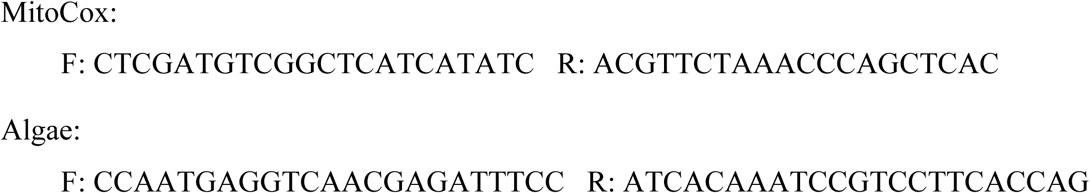

Five nanograms of DNA were used per reaction, derived from primary populations of *Oxytricha* FACS-derived nuclei, non-specific sorted “debris” particles, whole cell DNA, and lysates of single vegetative *Oxytricha* cells. Enrichment of targets was calculated as 2^(Cq[Sample]-Cq[Single^ ^Cell])^ with a minimum enrichment score of 0.

### Hi-C library preparation

Hi-C libraries were built from sorted nuclei largely as in Belaghzal et al.,^57^ starting from the post-cell lysis stage. 1-2 million sorted micronuclei were used as input for vegetative samples, while 250,000-500,000 anlagen were used as input for 24h developmental samples and 80,000 for 48h developmental Hi-C. Fixed, sorted nuclei were pelleted for 5 minutes at 4,000xg and resuspended in 342µl NEBuffer3.1 (New England Biolabs). 38µl %1 sodium dodecyl sulfate was added and samples were incubated for 10 minutes at 65°C. 43µl 10% Triton X-100 was then added. DNA was then digested with 400U DpnII (New England Biolabs) overnight at 37°C in a ThermoMixer with gentle shaking. The next day, the following reagents were added and samples were incubated at 23°C for 4h in a ThermoMixer with intermittent shaking: 1.5µl each of 10mM dCTP, dGTP, and dTTP (Thermo Fisher Scientific), 37.5µl 0.4mM biotin-14-dATP (Thermo Fisher Scientific), 6µl 10x NEBuffer 3.1, 2µl Dnase-free water (Thermo Fisher Scientific), and 10µl 5U/µl DNA polymerase I Klenow (New England Biolabs). Blunt end ligation was then performed at 16°C for 4h in a ThermoMixer with intermittent shaking with the following reagents: 24µl water, 240µl 5x ligation buffer (Invitrogen), 120µl 10% Triton X-100, µl 10 mg/ml BSA, and 50µl T4 DNA ligase (Invitrogen). Crosslinking was reversed with 50µl 10mg/ml proteinase K and a 2 hour incubation at 65°C, followed by addition of another 50µl proteinase K and overnight incubation at 65°C. The following day, DNA was purified by phenol-chloroform extraction, followed by incubated with 1µl 1mg/ml RnaseA (Thermo Fisher) for 30 minutes at 37°C. Biotin was removed from unligated ends by treating DNA with 5µl 10x NEBuffer 2.1 (New England Biolabs), 0.125µl 10mM dATP, 0.125µl 10mM dGTP, and 5µl 3,000 U/ml T4 DNA polymerase (New England Biolabs) in a total reaction volume of 50µl and incubating at 20°C for 4h followed by 75°C for 20 minutes. DNA was then washed and collected using a 30kD Amicon column (Millipore).

Illumina libraries were then prepared from Hi-C DNA using the NEBNext Ultra II kit (New England Biolabs). DNA was first sheared to 200-300bp using a Covaris M220 ultrasonicator (Covaris). After end preparation, and prior to adapter ligation and PCR, biotin-containing fragments were pulled down with MyOne Streptavidin C1 beads (Thermo Fisher Scientific). Streptavidin beads were twice washed with 400µl Tween wash buffer (5 mM Tris-HCl, pH 8.0; 0.5 mM EDTA, 1 M NaCl, 0.05% Tween 20), rocked for 3 minutes at room temperature, then reclaimed on a magnetic rack. 400µl 2x binding buffer (10 mM Tris-HCl, pH 8.0; 1 mM EDTA, 2 M NaCl) was added, along with the end prepped DNA and TLE buffer (10 mM Tris-HCl; 0.1 mM EDTA, pH 8.0) up to 800µl total. DNA was bound to beads for 15 minutes at room temperature with rotation, after which beads were reclaimed, washed first with 400µl binding buffer, then with 400µl TLE, before being resuspended in 20 µl TLE. The remaining steps of NEBNext Ultra II library prep were performed on-bead. After 3-8 cycles of indexing PCR, final libraries were purified using Ampure XP beads (Beckman Coulter).

Hi-C libraries were analyzed via Bioanalyzer and sequenced on NextSeq 500 instruments (Illumina) at the Columbia Genome Center. All Hi-C datasets were comprised of 2×75 and 2×150 paired-end reads. A summary of all Hi-C datasets is available in Table S1. Aliquots of the vegetative germline Hi-C library were set aside for quality control via forward-forward PCR. Three pairs of primers spanning DpnII GATC motifs and pointing in the same direction were designed, such that no PCR product is expected in the unprocessed germline genome. PCR was performed on unprocessed germline DNA and the vegetative germline Hi-C library to check for amplification exclusively from Hi-C chimeric DNA.

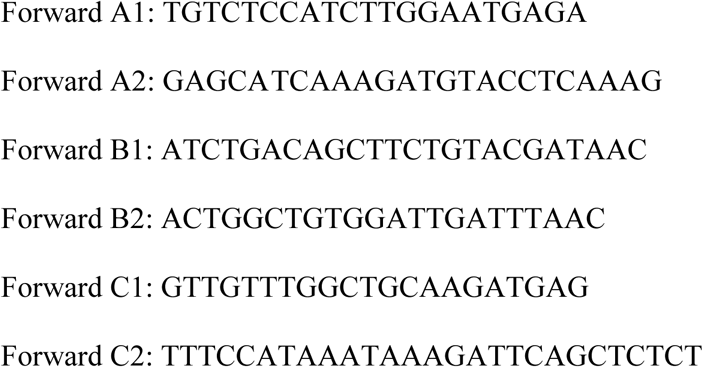

### RNase-treated Hi-C

For the RNase treated 24h Hi-C library, cells were pelleted prior to fixation by centrifugation for 2 minutes at 200xg. Cells were resuspended in 50µl 0.5x PBS with 0.02% Tween 20 and incubated on ice for 10 minutes. RNA was digested with 10µl 700U/ml RNase A (Thermo Fisher Scientific) for 30 minutes at room temperature in a Thermomixer at 950 rpm.

Under these low salt conditions (70 mM NaCl), RNase A cleaves single-and double-stranded RNA as well the RNA strand in RNA-DNA hybrids. For the control sample, 10µl 0.5x PBS was added rather than RNase A before incubation. Post digestion, samples were brought up to 50ml with Pringsheim media, pelleted, then washed and pelleted again in 50ml Pringsheim media before proceeding with the standard fixation, lysis, and sorting protocol as above. 10% of these samples were set aside for total RNA isolation using TRIzol (Thermo Fisher Scientific) and quantification of RNA by Qubit (Thermo Fisher Scientific).

### *Oxytricha* germline genome scaffolding

For Nanopore sequencing, DNA from sorted JRB310 *Oxytricha* micronuclei was isolated via phenol-chloroform extraction. The high molecular weight of the MIC DNA was confirmed on a 1% agarose gel. Two biological replicates of 450-550ng were prepared, and library preparation and long-read sequencing of these were performed at Oxford Nanopore Technologies New York lab. The combined Nanopore MIC dataset consisted of 13.8 Gb of long-read data. Reads with a GC content outside of the expected range for *Oxytricha* DNA (20-45%) were discarded. The remaining reads were used to scaffold the original draft germline genome assembly^40^ with LRScaf.^59^ Nanopore reads were first mapped to the draft assembly using Minimap2,^101^ and then LRScaf was run with the following settings: maximum overhang length 600, maximum overhang ratio 0.3, minimum overlap ratio 0.3. The LRScaf output was then fed into the pipeline for 3D-DNA.^60^ JRB310 vegetative MIC Hi-C reads were mapped to the LRScaf output using Juicer (v1.6).^58^ The subsequent merged_nodups.txt file was provided as input to 3D-DNA with the following settings, given the abundance of very short contigs in the draft assembly: size threshold 1000, polisher input size threshold 10000, splitter input size threshold 20000. The resulting.hic matrix and assembly were reviewed in Juicebox Assembly Tools^61^ to manually correct any suspected mis-joins. After the 3D-DNA assembly pipeline was completed, a final round of LRScaf was performed with the same settings as above. All resulting contigs less than 10 kb were checked against the nr database using BLAST,^62^ and contigs with exclusively prokaryotic hits in the top 10 reported results were removed from the scaffolded assembly. MDS and TBE annotations from the initial fragmented assembly were mapped onto the scaffolded assembly via BLAST, requiring 98% identity to the queries.

### Hi-C data processing

Hi-C Illumina reads were assessed using FastQC^102^ and were trimmed using Trimmomatic.^103^ Reads were then mapped to the scaffolded genome assembly using the Juicer (v1.6) pipeline.^58^ For data visualization and subsequent analyses, Juicer output was converted to. h5 matrices at resolutions of 1000, 5000, 10000, and 100000 using HiCExplorer (v3.7.1).^31,67,68^ PCR duplicate removal, KR matrix balancing^63^ and normalization between samples was performed within HiCExplorer, as were PCA,TAD, and log2 analyses at various resolutions.

Raw matrix counts were exported for analysis of interactions between specific bins. Compartments were assigned on the basis of the sign of the first eigenvector following PCA of observed/expected Hi-C matrices. TAD calling was performed using the hicFindTADs function at resolutions of 5000, 10000, and 100000, using f.d.r. correction for multiple testing, minDepth of 3x[resolution], maxDepth 10x[resolution], step equal to [resolution], and delta threshold of 0.05. Overlapping TAD calls from multiple resolutions were merged using the hicMergeDomains function.

### ATAC-seq

ATAC-seq (Assay for Transposase-Accessible Chromatin sequencing) was performed on fixed, sorted nuclei following previously published protocols.^104^ For vegetative samples, at least 500,000 fixed and sorted micronuclei were used as input for each sample. For developmental samples, at least 100,000 24h anlagen were used as input. Three replicate libraries of each were prepared with varying amounts of Tn5 transposase (Illumina). Sorted nuclei were pelleted for 5 minutes at 4,000xg and resuspended in 50µl lysis buffer (10mM Tris–HCl, pH 7.4; 10mM NaCl; 3mM MgCl_2_; 0.05% Igepal CA-630), then centrifuged again at 4,000xg for 5 minutes. Nuclei pellets were resuspended in 50µl 1x Tagment DNA Buffer (Illumina) along with, for each type of input nucleus, 1µl, 2.5µl, or 5µl Tn5 transposase (TDE1 enzyme, Illumina). Transposase reactions were incubated for 30 minutes at 37°C. After transposition, a reverse-crosslinking solution was added up to 200µl, with a final concentration of 50mM Tris–HCl, 1mM EDTA, 1% SDS, 0.2M NaCl, and 5 ng/ml proteinase K, and samples were incubated overnight at 65°C in a Thermomixer at 1,000 rpm. The next day, DNA was extracted via NucleoSpin Mini kit (Macherey-Nagel), and Illumina libraries were prepared with NEBNext Ultra II. Libraries were sequenced on a NextSeq 500 at the Columbia Genome Center, generating between 10 million and 13.5 million 2×75 paired-end reads per sample. The quality of reads was assessed using FastQC^102^ and reads were trimmed using Trimmomatic.^103^ Reads were mapped to the scaffolded germline assembly using Bowtie2^105^ in “very-sensitive” mode. For peak and coverage analyses, replicates treated with different quantities of Tn5 were merged. Narrow peaks were called from mapped ATAC-seq reads using MACS2.^106^ Average coverage along MDSs, IESs, and TBEs was assessed by identifying features of a narrow size range (500-1,000bp for MDSs and IESs, 3-4 kb for TBEs), then building coverage matrices and plotting coverage profiles using Deeptools (v3.5.1),^107^ with 500 bp upstream and downstream of features included.

### DNA fluorescence *in situ* hybridization

Potential representative 40 kb regions for compartments A and B were identified on the basis of PC1 scores, and probes ranging from 35-40 nucleotides were designed to cover these regions using OligoMiner^72^ and BLAST.^62^ Three regions on which to perform FISH from each compartment were then selected on the basis of available unique sequences, in order to have sufficient specific probe coverage, with a target of 8 probes per kilobase. OligoFISH probe pools were then ordered from and manufactured by Daicel Arbor Biosciences. For each compartment, one probe pool was labeled with Alexa 488, a second pool was labeled with ATTO 550, and a third probe pool was labeled with ATTO 647N. Probe pools were stored at-20°C at a concentration of 10 pmol/µl in TE buffer (10 mM Tris,1 mM EDTA, pH 8.0).

*Oxytricha* cells on which FISH was performed were prepared by mating JRB310 and JRB510 mating types, harvesting at 24h, and fixing for 10 minutes in a 1% formaldehyde (v/v) solution in Pringsheim at room temperature with gentle shaking. Glycine was then added to a final concentration of 125mM to quench fixation, followed by incubation at 4°C for 15 minutes with gentle shaking. Cells were pelleted for 3 minutes at 300xg, washed once with 1xPBS, then pelleted again and stored in 1xPBS at 4°C.

A protocol for 3D-FISH in *Oxytricha* was developed based on previous recommendations for other systems.^71^ 24h developmental *Oxytricha* cells were affixed overnight at 4°C to slides coated in poly-L-lysine (Sigma-Aldrich). Cells were washed with 20µl 1X PBS for 1 minute at room temperature. Cells were then permeabilized by treatment with 0.5% Triton X-100 (v/v) in 1X PBS for 30 minutes at room temperature, followed by washing with 1X PBS for 2 minutes at room temperature, and then treatment with 0.1N HCl for 5 minutes at room temperature. Cells were then washed twice with 2X saline-sodium citrate buffer (SSC) (Thermo Fisher Scientific) for two minutes each at room temperature. Pre-hybridization was performed by equilibrating cells for 5 minutes at room temperature in 40% formamide in 2X SSC, followed by treating with 40% formamide in 2X SSC for 20 minutes at 60°C. Meanwhile, pools of probes in hybridization buffer were prepared. For cells probed for compartment A, 1.5 pmol of each compartment A probe pool were used; for cells probed for compartment B, 1.5 pmol of each compartment B probe pool were used. Probes were added to 20µl hybridization buffer consisting of final concentrations of 40% formamide (v/v), 2X SSC, 0.5U/µl RNase A (Thermo Fisher Scientific), and 10% dextran. The mixture of probes in hybridization buffer was pre-heated at 70°C for 5 minutes and then added to equilibrated cells. Slides were transferred to a humid container, then denatured for 2.5 minutes at 78°C before being incubated at 42°C overnight.

After overnight hybridization, slides were mounted with VECTASHIELD Antifade Mounting Medium with DAPI (Vector Laboratories). Slides were imaged with a Nikon Ti2 AXR MP confocal microscope at 100X with 0.5µm Z steps. Images were processed and single Z-slices were selected using Fiji.^108^

### 3D reconstructions of scaffolds guided by the empirically measured pairwise *cis*-interactions, captured by Hi-C experiments

The 3D reconstructions were produced using a modified 4DHiC method,^73^ which enables interactions between multiple chromosomes. Here, an extended, disordered beads-on-springs polymer chain is collapsed using additional constraints defined by the empirically determined

Hi-C contact map, using contacts exceeding a threshold and separate levels of constraints are used for intra-and inter-chromosomal contacts.^74^ Specifically, all simulations were conducted using the Large-Scale Atomic/Molecular Massively Parallel Simulator (LAMMPS).^75^ Each scaffold consists of several coarse-grained (CG) beads, with each bead representing 20 kb resolution. The total number of CG beads varies depending on the length of the individual scaffold. For example, Scaffold 528 (5.06 Mb) was represented by a polymer with 253 coarse-grained (CG) beads. The initial structure is modeled as a connected chain, which was generated through a self-avoiding random walk algorithm. The pair interactions between the CG beads were simulated using the FENE and Lennard-Jones potentials. Experimental Hi-C constraints were applied as harmonic constraints to simulate cross-linking contacts. To minimize noise from low-intensity signals, the frequency of Hi-C contacts was set to be greater than 10 for all intra-and 20 for all inter-scaffold interaction, respectively. The simulation temperature was maintained at a constant value using the Langevin thermostat, set at 1 in reduced units, with a damping coefficient of 1 *τ*⁻¹. A timestep of 0.01 *τ* was utilized, where τ represents the reduced (Lennard-Jones) time. The polymer, subjected to harmonic constraints, was analyzed using Brownian dynamics under implicit solvent conditions. To compare the similarities between the experimental and simulated Hi-C maps, the Pearson correlation coefficient was calculated. The built-in LAMMPS routines were employed to calculate the radius of gyration (*R_g_*). To understand the spatial organization of compartments in 3D, the experimental A and B compartment PC_1_ scores were mapped onto simulated 3D models.^76,77^

### DNA-RNA immunoprecipitation

DNA-RNA hybrids were captured from whole cell DNA using the S9.6 antibody (Sigma-Aldrich). DNA was purified via phenol-chloroform extraction from a mating timecourse, with samples taken every 12 hours. Half the extracted DNA was digested with 15µl 5U/µl RNaseH in 1x Reaction Buffer (Thermo Fisher Scientific) for 2h at 37°C and then purified again by phenol-chloroform. 130ng DNA was loaded onto a nitrocellulose membrane for each timepoint and each condition (+/-RNaseH). The membrane was blocked for 2.5h while rocking in TBS-T (0.05% Tween-20, 20 mm Tris-HCl, 150 mm NaCl, pH 7.5) with 5% BSA (w/v). The membrane was then incubated with the S9.6 antibody for 1 hour at a 1:2,000 dilution in TBS-T with 0.1% BSA (w/v), then washed three times in TBS-T for 5 minutes each, followed by incubation with secondary goat anti-mouse horseradish peroxidase-conjugated secondary antibody diluted 1:1,000 in TBS-T with 5% BSA for 30 minutes. The membrane was washed for 15 minutes in TBS-T and then for 5 minutes in TBS (20 mm Tris-HCl, 150 mm NaCl, pH 7.5), after which it was placed within a plastic sheet, Pierce ECL substrate (Thermo Fisher Scientific) mixture was added, and the membrane was kept out of light for 1 minute before being imaged via chemiluminescence on an Amersham Imager 600 (GE).

For DRIP-seq library prep, genomic DNA from approximately 500,000 cells each was extracted from a mating timecourse every 12h via phenol-chloroform up until 60h.

Immunoprecipitation generally followed the protocol in Wahba et al.^88^ Half of each sample was used for RNaseH-treated controls, which were incubated with 10µl 5U/µl RNaseH in 1x Reaction Buffer (Thermo Fisher Scientific) overnight at 37°C before being repurified by NucleoSpin (Macherey-Nagel). DNA from RNaseH-treated and-untreated samples was then sonicated in an S220 Covaris machine to 350bp. For each sample, 5µg of S9.6 antibody was bound to 50µl Protein A Dynabeads (Thermo Fisher Scientific) for 1 hour at 4°C. Beads were recovered on a magnetic rack, and sonicated DNA samples were then incubated with S9.6-bound Dynabeads in 100µl 1X FA (1% Triton X-100, 0.1% sodium deoxycholate, 0.1% SDS, 50mM HEPES, 150mM NaCl, 1mM EDTA) for 2h rotating at 4°C. Beads were then recovered and serially washed once with 1X FA, once with 1X FA with 0.5M NaCl, once with lithium chloride detergent solution (0.25M LiCl, 0.5% NP-40, 0.5% sodium deoxycholate, 1mM EDTA, and 10mM Tris-HCl), and twice with TE buffer. DNA was eluted from the beads with 300ul elution buffer (1% SDS, 0.1M NaHCO_3_). After elution, 5µl RNase Cocktail (Thermo Fisher Scientific) in 50µl 0.5X PBS was added to the beads, followed by rotation at room temperature for 15 minutes, after which that supernatant was reclaimed and then added to the initial eluate. Samples were then treated with 1µl 20mg/ml proteinase K (Takara Bio) for 1 hour at 42°C in a Thermomixer at 1000 rpm. DNA was then purified via phenol-chloroform extraction, and Illumina libraries were prepared with NEBNext Ultra II. Libraries were sequenced on a NextSeq 500 at the Columbia Genome Center, yielding at least 4.5 million 1×75 single-end reads per sample. The quality of reads was assessed using FastQC^102^ and were trimmed using Trimmomatic,^103^ before being mapped to the *Oxytricha* somatic reference genome^38^ using BWA MEM.^109^ 5,000 somatic contigs were identified between 1.5 kb and 3 kb and bearing single genes on the same strand, and DRIP samples at 0h and 24h were normalized to their RNaseH-digested controls using the difference operation in Deeptools bamCompare, after which the resulting matrices were averaged over the selected somatic contigs and plotted in Deeptools.^107^

### Quantification and statistical analysis

All statistical tests were performed in R (v4.1.2) and described in figure legends.

