## Supplemental material for "Widespread 3D genome reorganization precedes programmed DNA rearrangement in *Oxytricha trifallax*"

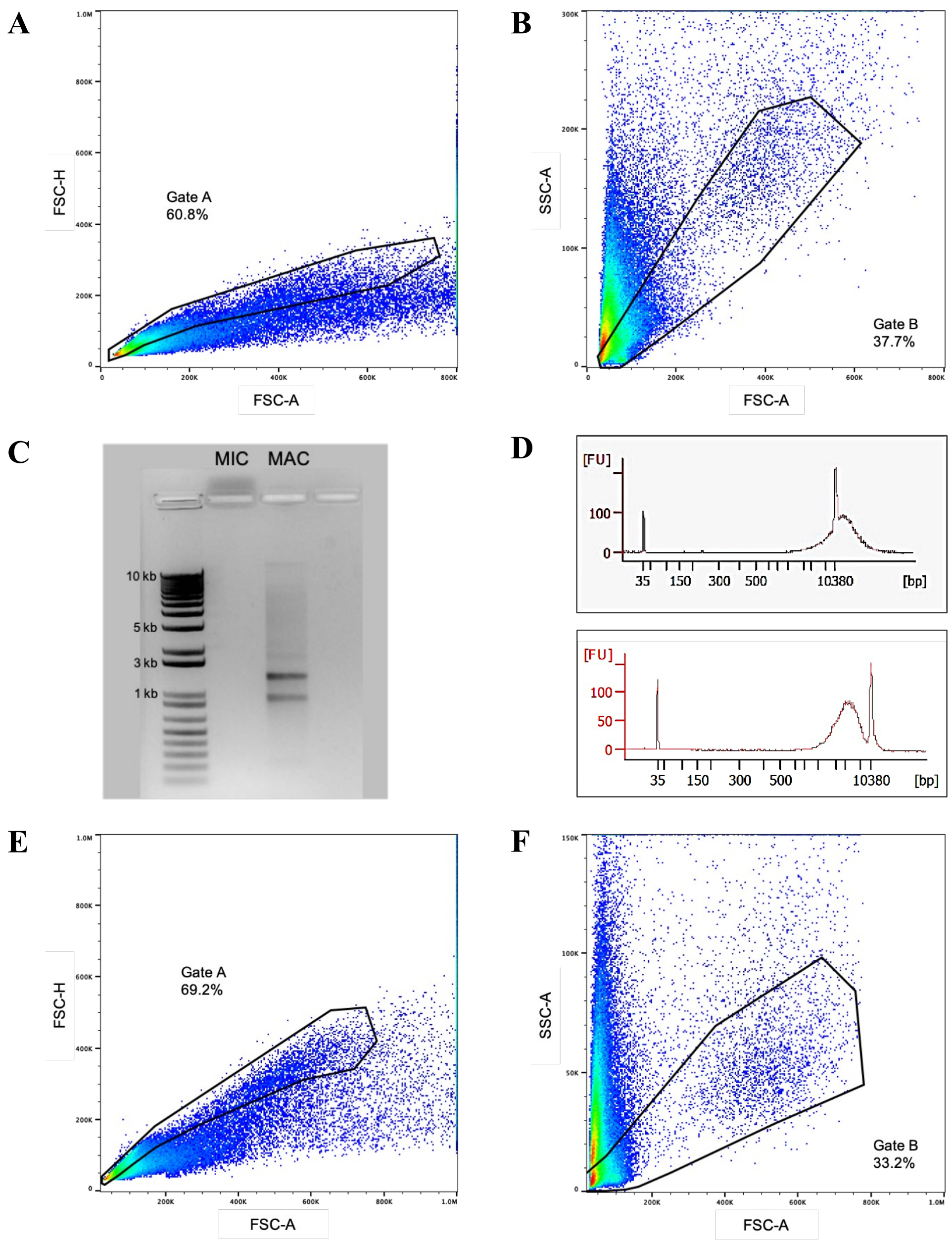


**Figure S1.** Gating of FACS Samples.
(A) Forward scatter area x forward scatter height of vegetative nuclei.
(B) Forward scatter area x side scatter ("SSC") area of vegetative nuclei.
(C) 1.5% agarose gel of DNA derived from vegetative MIC and MAC FACS populations.
(D) Bioanalyzer High Sensitivity trace of DNA from sorted vegetative MIC (top) and MAC (bottom).
(E) Forward scatter area x forward scatter height of 24hr developmental nuclei.
(F) Forward scatter area x side scatter area of 24hr developmental nuclei.

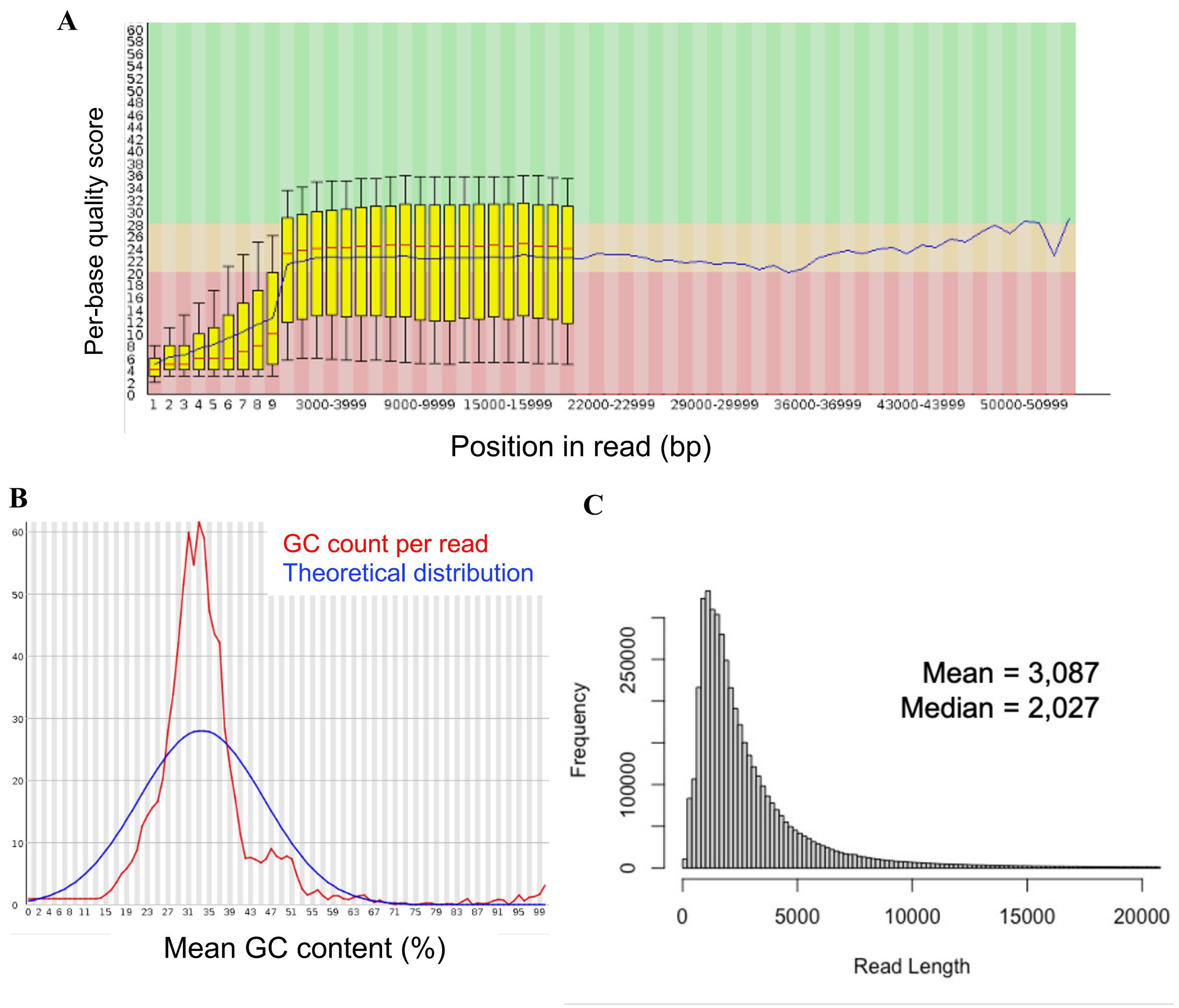


**Figure S2.** Quality metrics of Nanopore data.
(A) Average per-base quality score of subset of 10,000 reads (from FastQC[102]). Low quality reads were discarded.
(B) Average GC content of subset of 10,000 reads (from FastQC[102]). Reads with GC content outside of expected range for *Oxytricha* were discarded.
(C) Histogram of Nanopore read length. ~1% of reads out of graph range, up to maximum of 55 kb.

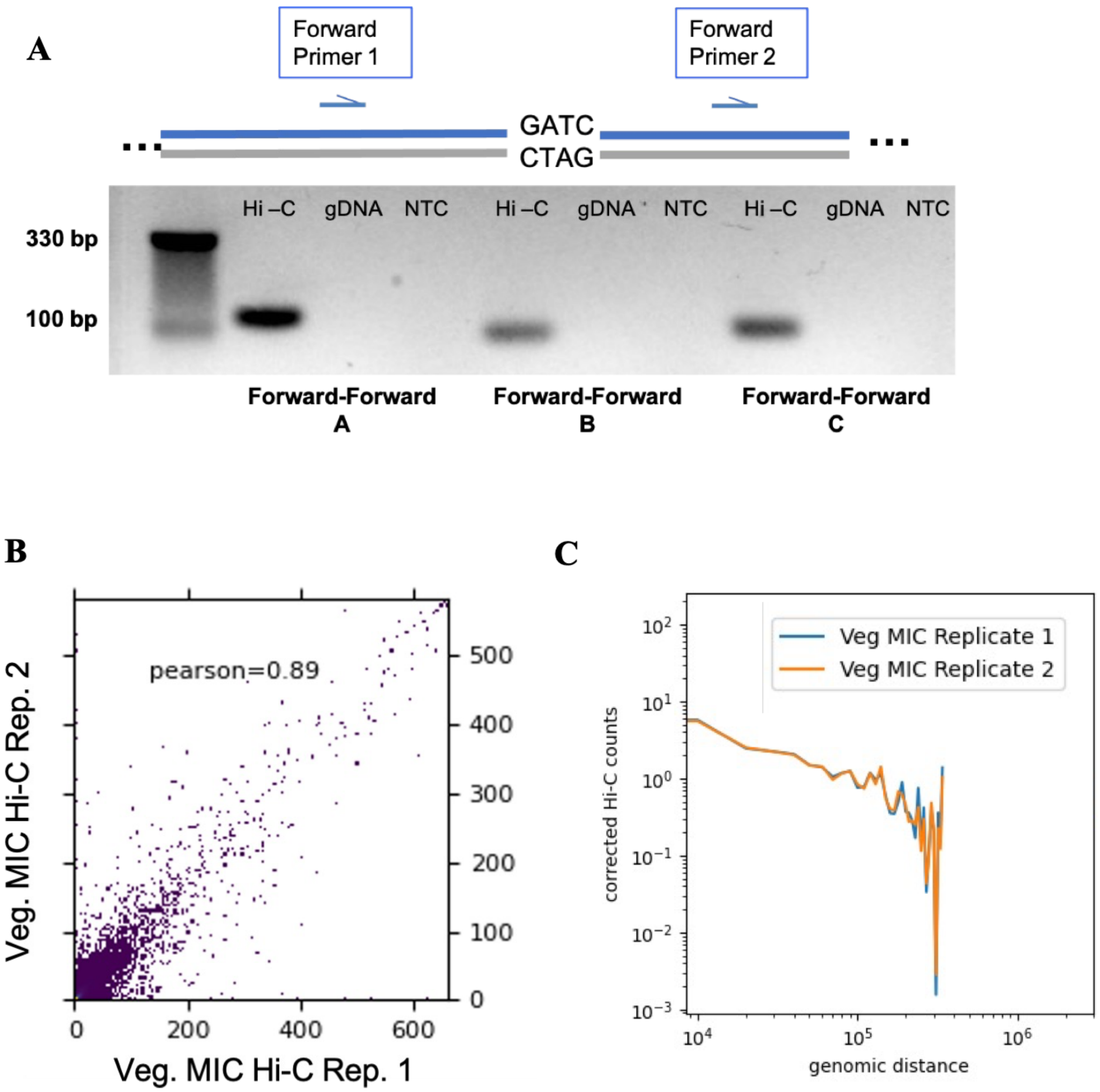


**Figure S3.** Quality control of vegetative germline Hi-C replicates.
(A) Forward-forward PCRs of vegetative MIC Hi-C libraries and genomic DNA.
(B) Correlation between vegetative MIC Hi-C replicates mapped to original reference (10 kb bins).
(C) Frequency of intrachromosomal Hi-C contacts across genomic distances in MIC Hi-C replicates mapped to original reference genome.

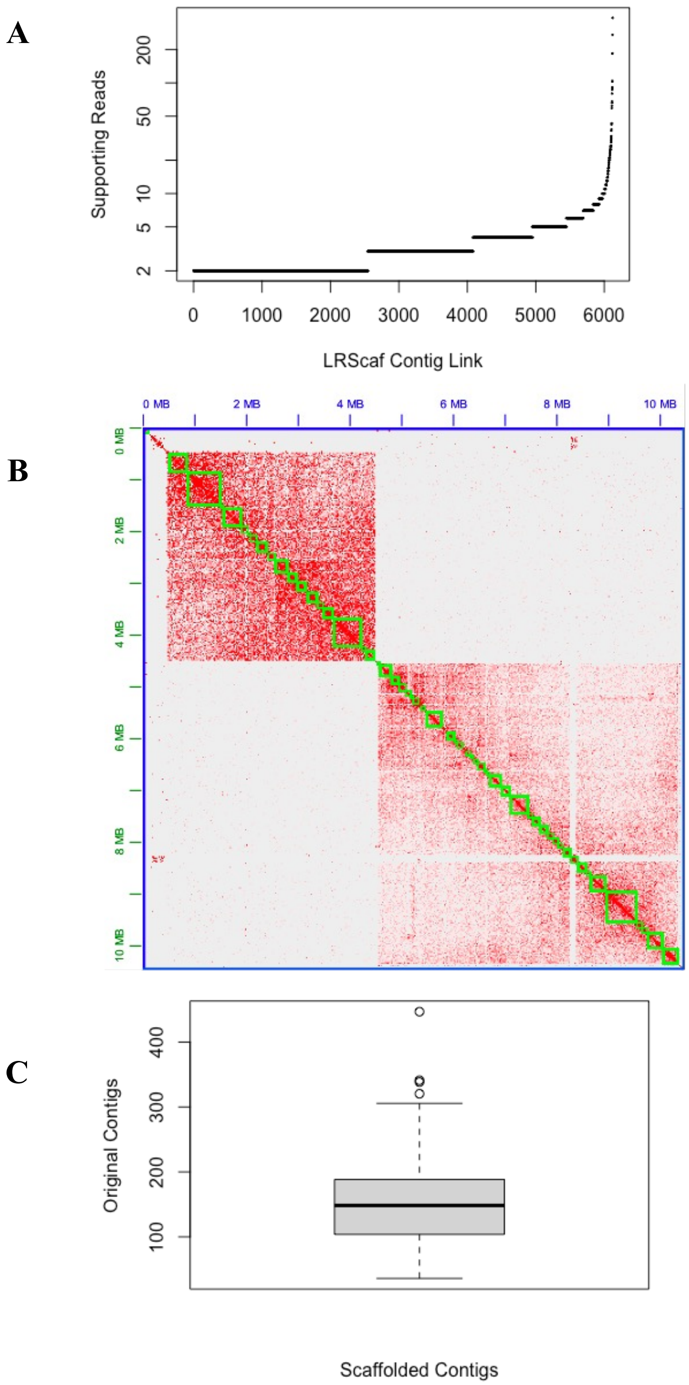


**Figure S4.** Methodology of genome scaffolding.
(A) Number of Nanopore reads supporting each LRScaf contig linking event.
(B) Manual review in Juicebox[61] of vegetative germline Hi-C contact frequency for two large 3D-DNA-derived scaffolds. Green boxes are contigs from original assembly.
(C) Number of contigs from original assembly that map to scaffolds (>1Mb).

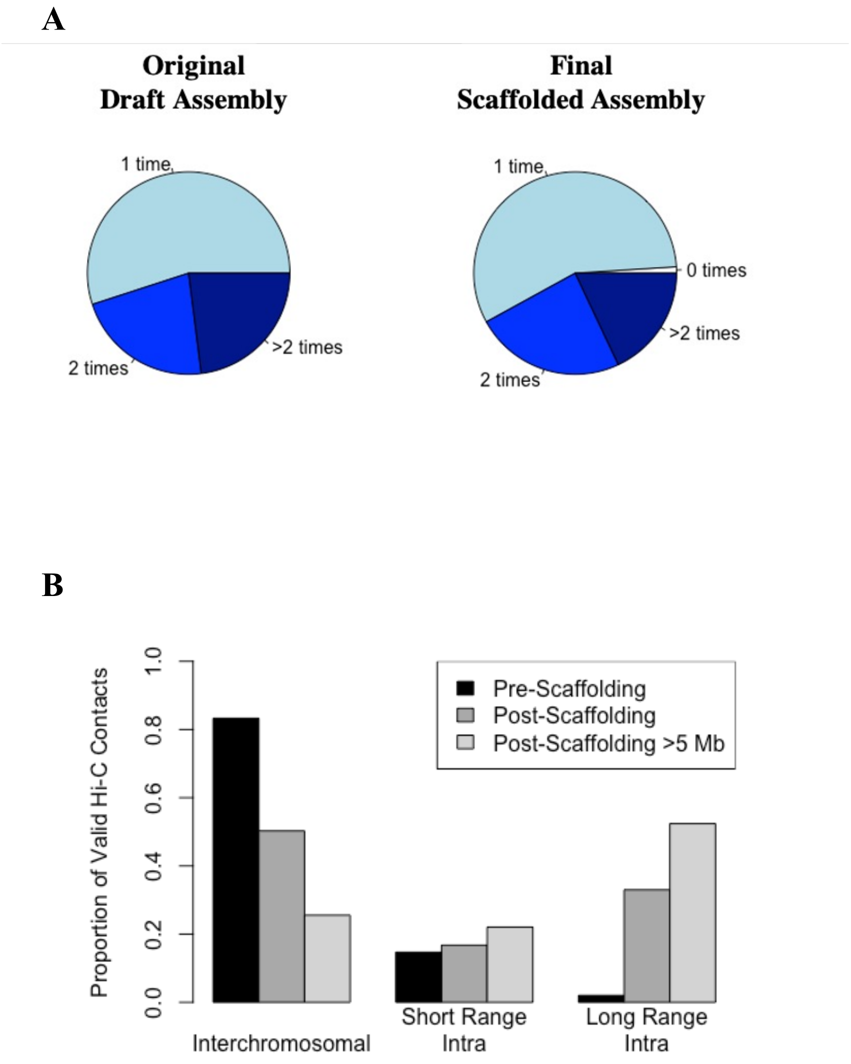


**Figure S5.** Comparison of data mappability between original and scaffolded assemblies.
(A) Number of BLAST hits for each somatic MDS queried against original or scaffolded assembly.
(B) Frequency of vegetative germline Hi-C contacts when mapped to original assembly and to scaffolded assembly, and the subset of the latter involving the largest scaffolds.

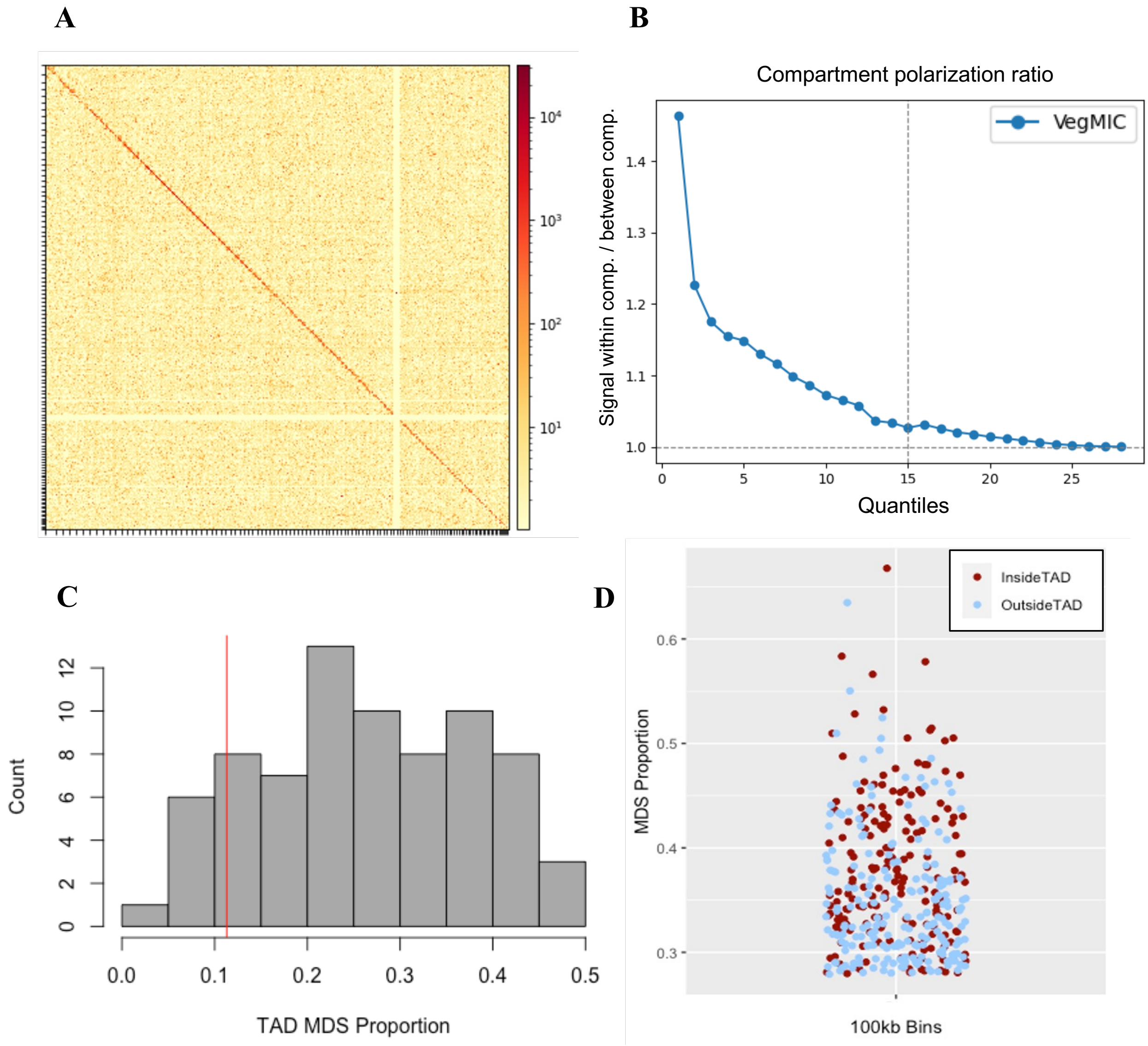


**Figure S6.** Vegetative germline shows high-order structure around the densest clusters of MDSs.
(A) Full genome-wide view of vegetative germline Hi-C data (log1p).
(B) Compartmentalization signal of the vegetative germline.
(C) Histogram of MDS content of TAD-like structures. Red line represents the genome-wide MDS content.
(D) Presence within TAD-like structures of the 10% most MDS-rich 100 kb bins in the germline.

**A**

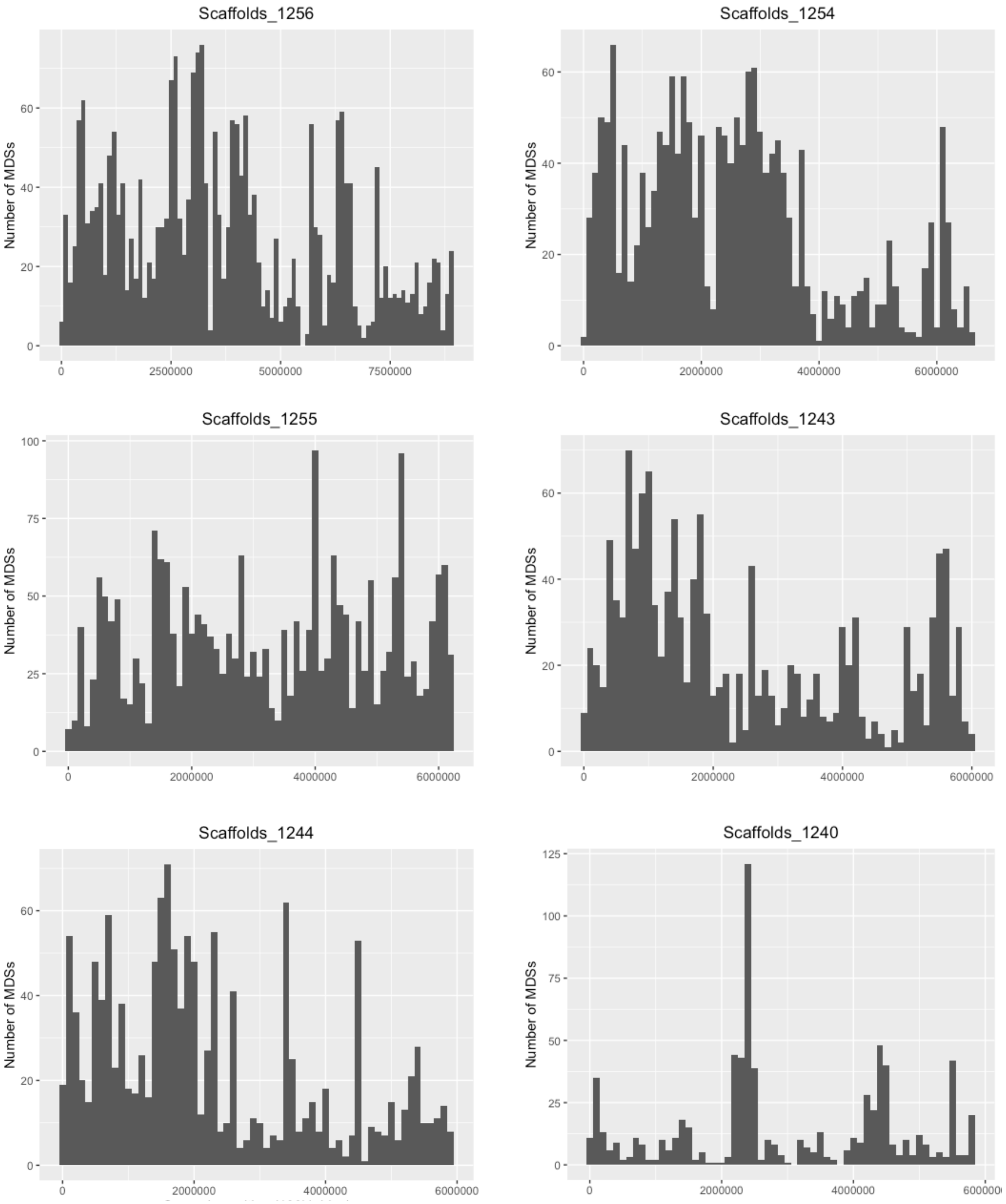


**Figure S7.** MDSs are distributed unevenly throughout the germline genome.
(A) MDS density along the 6 largest germline scaffolds.

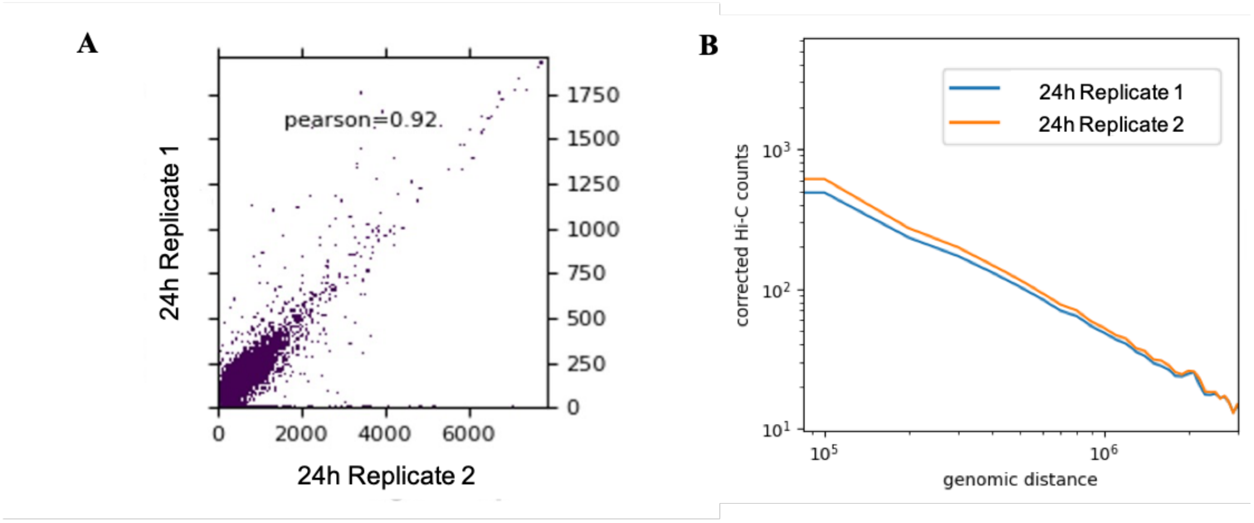


**Figure S8.** 24 hour anlagen Hi-C is highly replicable.
(A) Correlation of Hi-C contacts between 24h anlagen replicates (10 kb bins).
(B) 24h anlagen Hi-C replicates have similar distribution of intrachromosomal contact frequency vs. genomic distance.

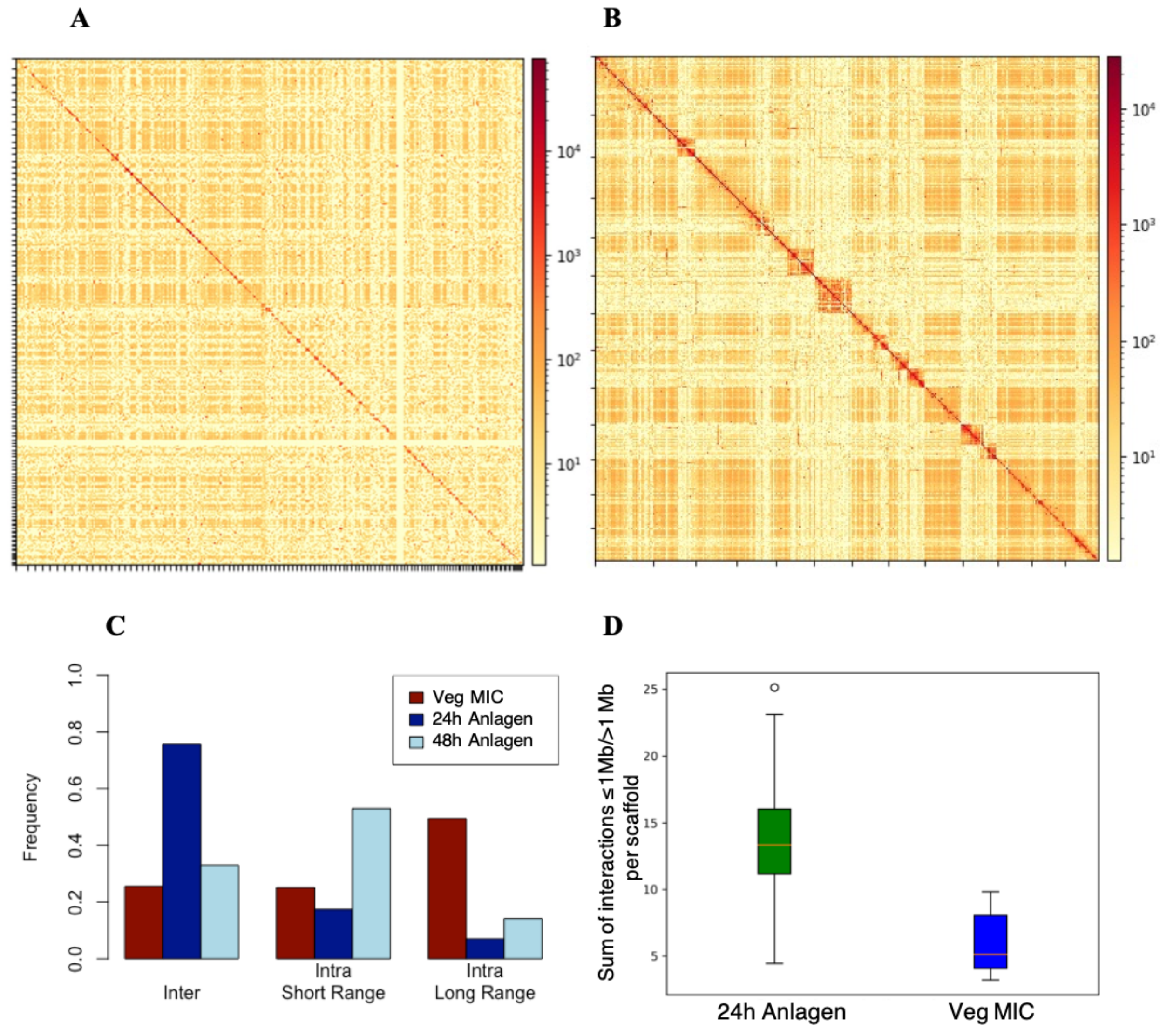


**Figure S9**. The developmental genome drastically reorganizes before rearrangement.
(A) Full genome-wide view of 24h anlagen Hi-C data (log1p contact frequency, 100 kb bins).
(B) 24h anlagen Hi-C data, scaffolds ≥ 5 Mb (log1p contact frequency, 100 kb bins).
(C) Frequency of different interaction types in vegetative germline, 24h, and 48h anlagen among scaffolds ≥ 5 Mb. Short-range is considered ≤ 20 kb.
(D) Distribution of ratio of intrachromosomal interactions ≤ 1Mb and > 1 Mb per scaffold, 24h anlagen vs. vegetative precursor germline.

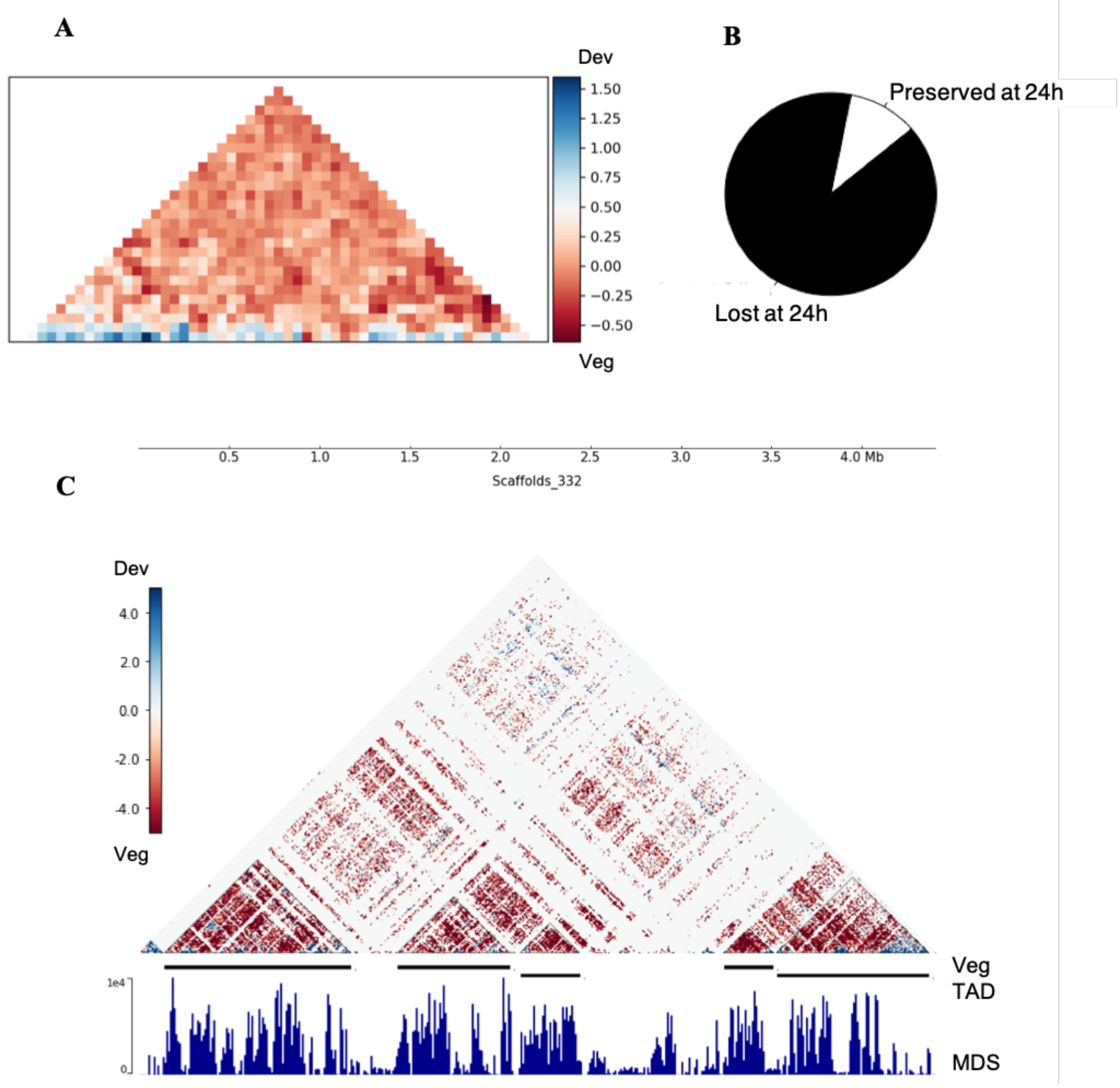


**Figure S10.** Vegetative germline TADs lose their structure by 24 hours.
(A) $log_2$ ratios between 24h anlagen and vegetative germline Hi-C averaged over all vegetative TAD-like structures.
(B) Proportion of vegetative TAD-like structures able to be called in 24h anlagen.
(C) 24h anlagen vs. vegetative germline $log_2$ ratios on scaffold shown in Figure 3 with several vegetative TAD-like structures.

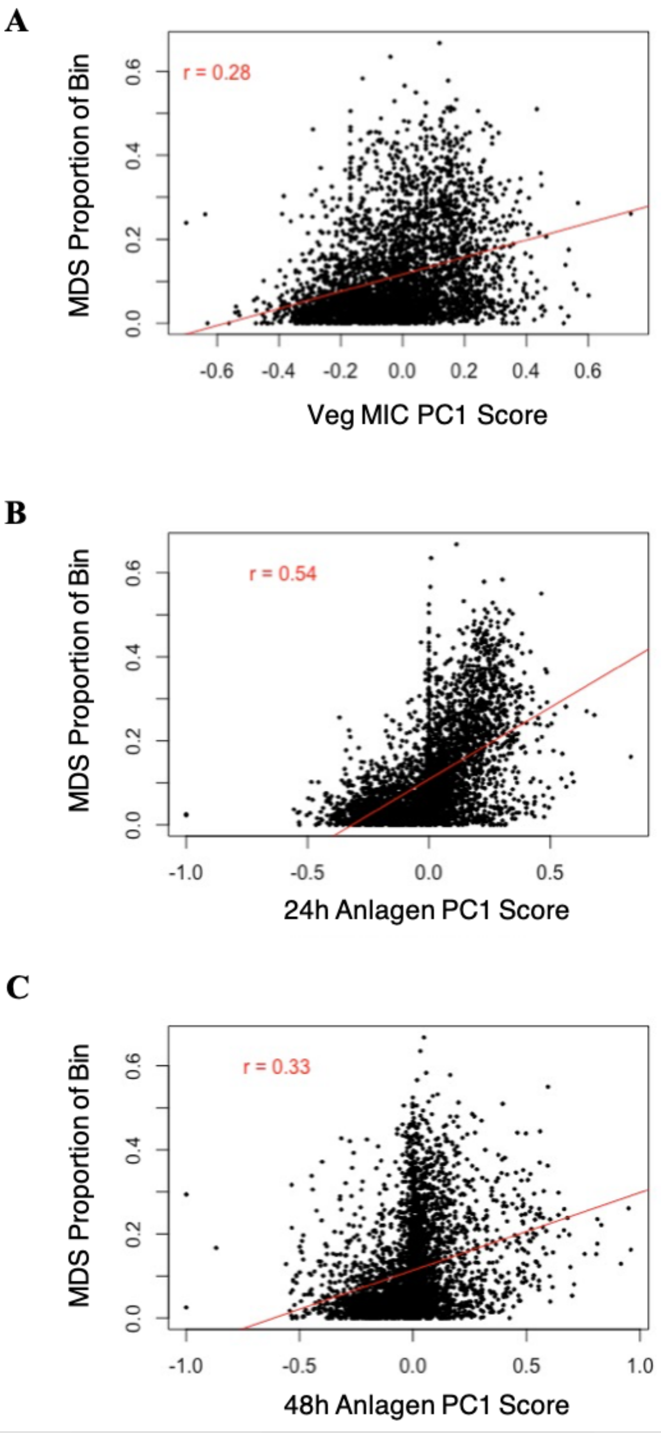


**Figure S11.** MDS proportion correlates with developmental compartment.
(A) Correlation in vegetative germline Hi-C of PCA1 value with MDS content, 100 kb bins. r is Pearson correlation coefficient.
(B) Same as above in 24h anlagen Hi-C.
(C) Same as above in 48h anlagen Hi-C.

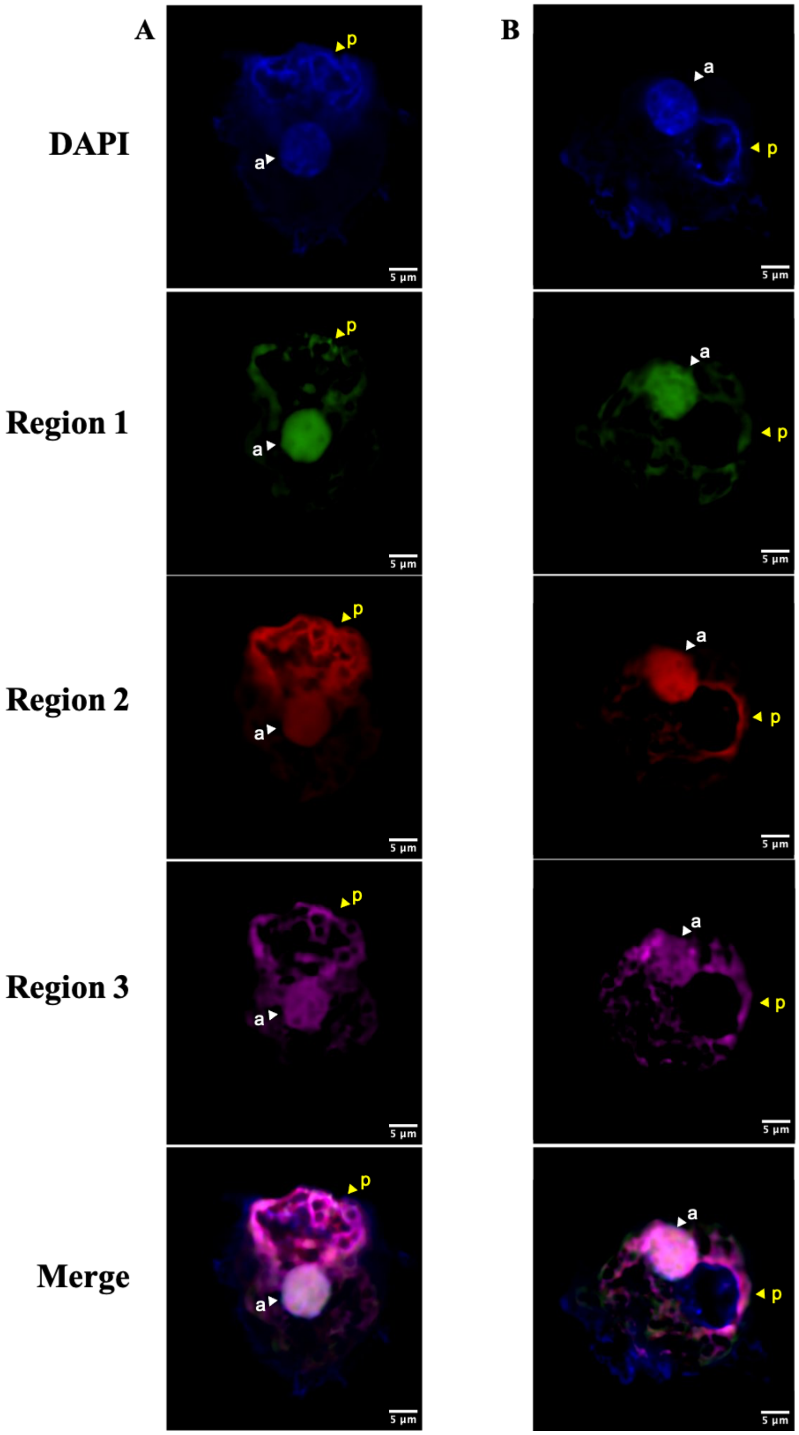


**Figure S12.** DNA FISH shows nonuniform distribution of compartmentalized DNA in the 24h anlagen.

(A) Single Z-slice of 24h cell probed for three non-syntenic 40kb regions in compartment A. (B) Single Z-slice of 24h cell probed for three non-syntenic 40kb regions in compartment B. “a” indicates developing nucleus; “p” indicates parental MAC.

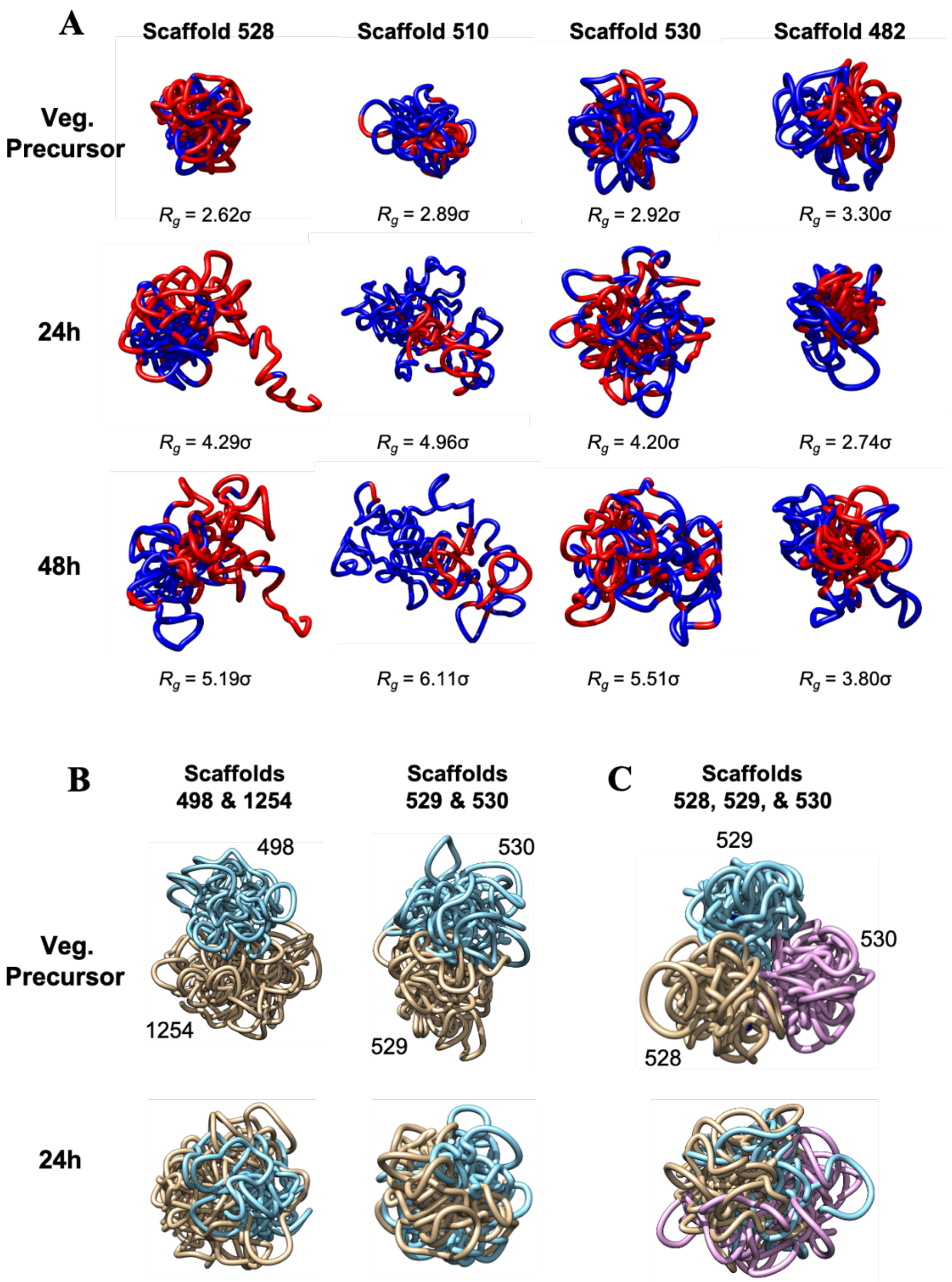
A
Scaffold 528
Scaffold 510
Scaffold 530
Scaffold 482
Veg. Precursor
$R_g$ = 2.62σ
$R_g$ = 2.89σ
$R_g$ = 2.92σ
$R_g$ = 3.30σ
24h
$R_g$ = 4.29σ
$R_g$ = 4.96σ
$R_g$ = 4.20σ
$R_g$ = 2.74σ
48h
$R_g$ = 5.19σ
$R_g$ = 6.11σ
$R_g$ = 5.51σ
$R_g$ = 3.80σ
B
Scaffolds 498 & 1254
Scaffolds 529 & 530
C
Scaffolds 528, 529, & 530
Veg. Precursor
498
1254
530
529
529
530
528
24h

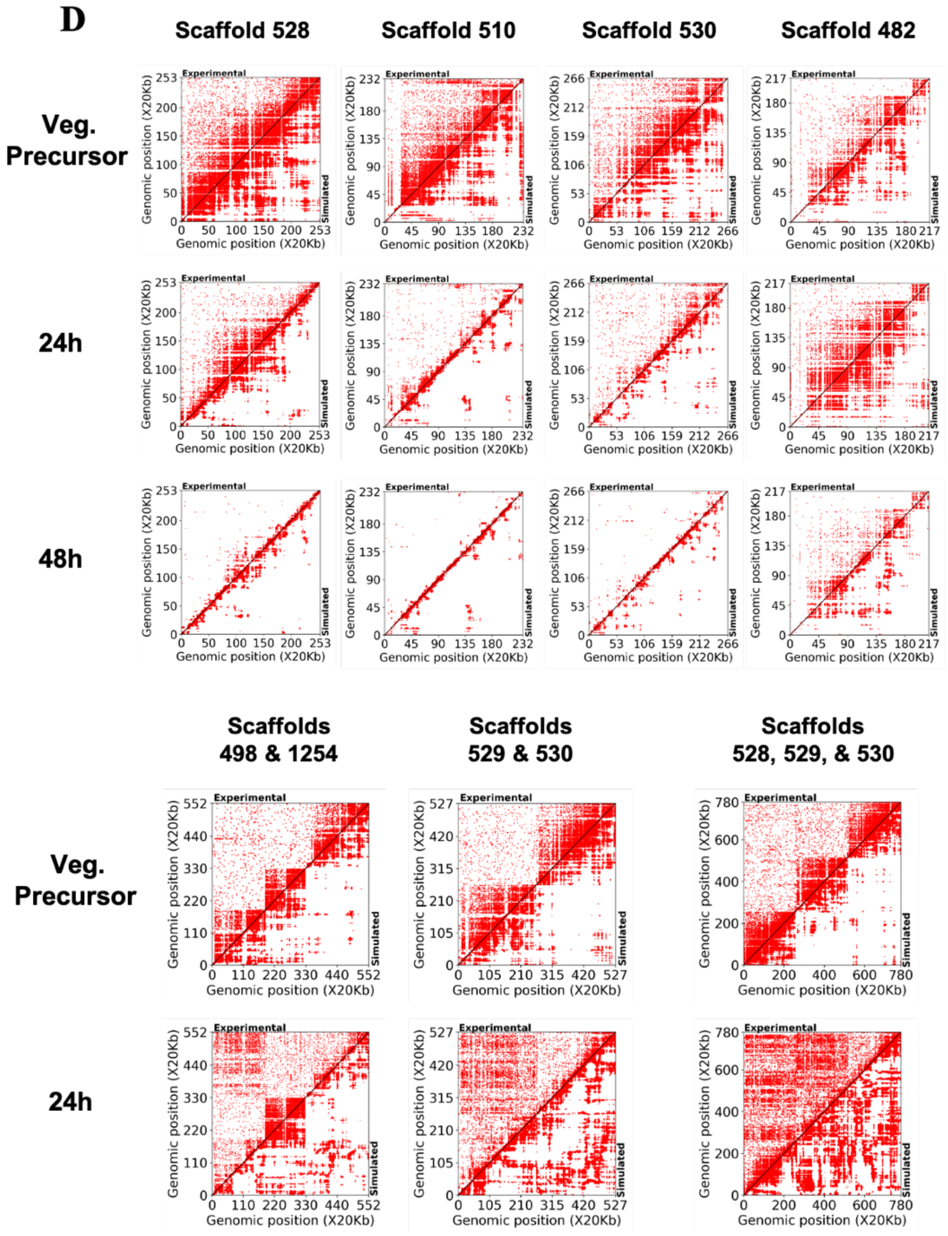


**Figure S13.** 3D reconstructions (guided by the empirically measured pairwise *cis*-interactions, captured by Hi-C experiments) illustrate significant reorganization of germline chromosomes during development.

(A) Single-scaffold 3D reconstructions of Hi-C data in the vegetative precursor and 24h and 48h developmental timepoints. Red indicates 20kb bead in compartment A; blue indicates 20kb bead in compartment B. All timepoints use 24h compartment data as basis for coloring. $R_g$ indicates radius of gyration.

(B) Two-scaffold 3D reconstructions. Coloring indicates separate scaffolds.

(C) Three-scaffold 3D reconstructions. Coloring indicates separate scaffolds.

(D) Comparisons of experimental (upper triangles) and simulated contact maps (lower triangles) show close agreement between experiments and simulations. Simulated contact maps are produced from 3D constructions shown in (A)-(C).

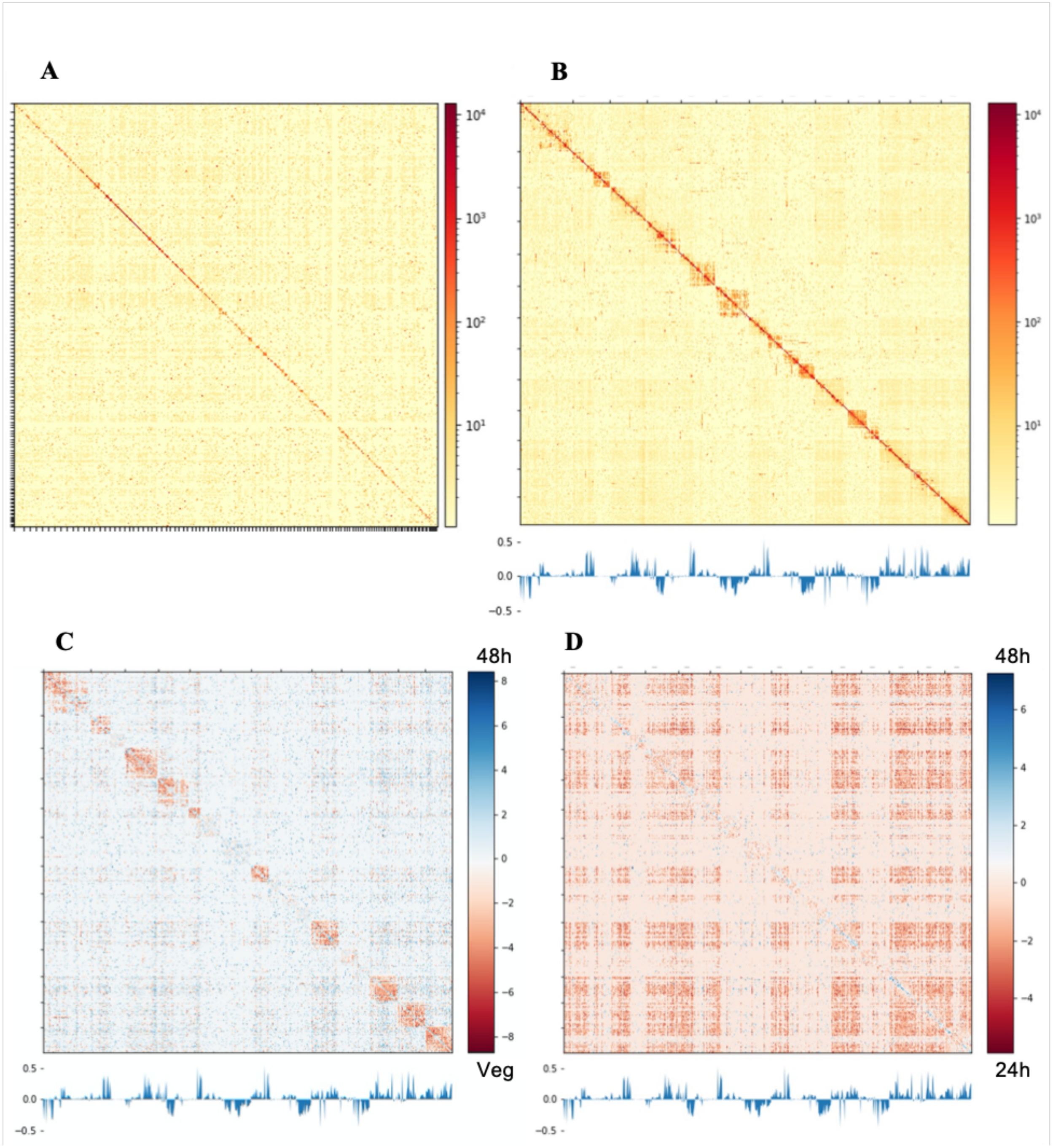


**Figure S14.** 48 hour anlagen shares reorganized structure with 24 hour anlagen but local contacts predominate.
(A) Genome-wide 48h anlagen Hi-C data, 100 kb bins, log1p.
(B) 48h anlagen Hi-C data across scaffolds ≥ 5 Mb, 100 kb bins, log1p.
(C) 48h anlagen vs. vegetative germline Hi-C $\log_2$ ratios across scaffolds ≥ 5 Mb, 100 kb bins.
(D) 48h anlagen vs. 24h anlagen Hi-C $\log_2$ ratios across scaffolds ≥ 5 Mb, 100 kb bins.

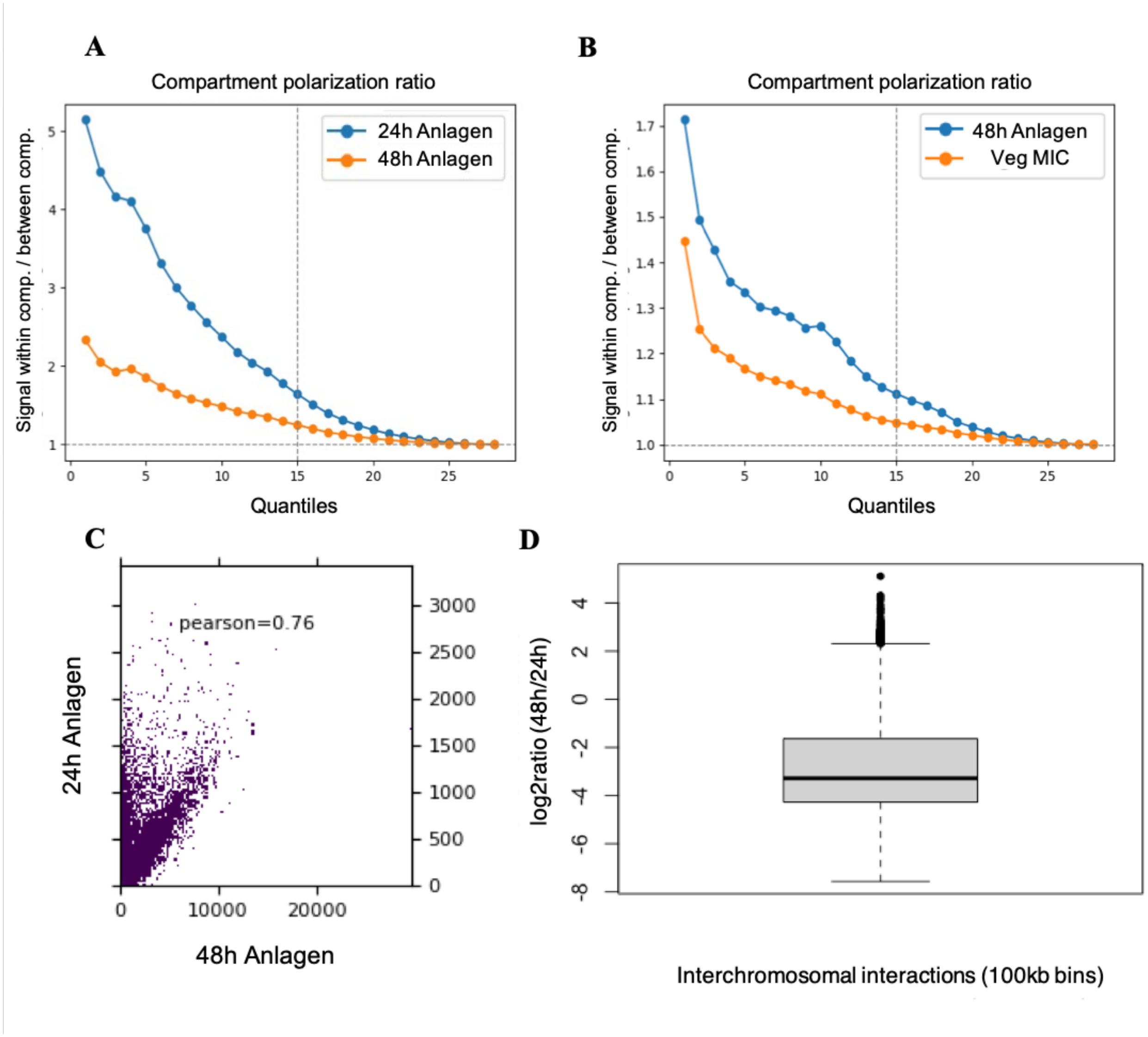


**Figure S15.** 48 hour anlagen loses most interchromosomal contacts.
(A) Compartmentalization scores of 24h vs. 48h anlagen (24h PCA1 used for building quantiles).
(B) Compartmentalization scores of vegetative germline vs. 48h anlagen (48h PCA1 used for building quantiles).
(C) Correlation of Hi-C matrices for 24h anlagen and 48h anlagen (10 kb bins).
(D) Comparison of interchromosomal interaction frequency in 48h anlagen vs. 24h anlagen, within 100 kb bins of germline scaffolds ≥ 5 Mb.

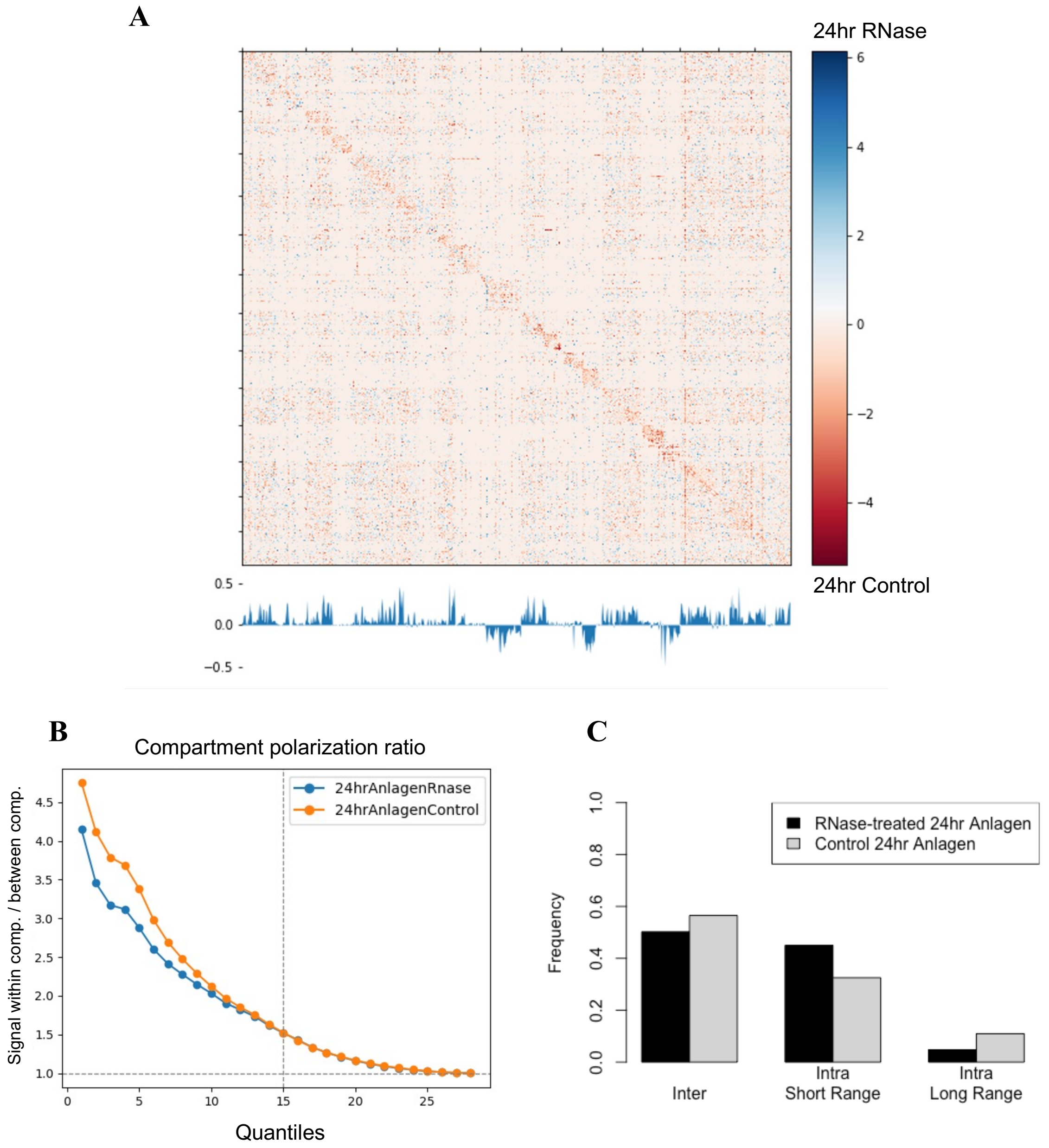


**Figure S16**. RNase treatment affects long-range contact frequency but not compartmentalization. (A) $Log_2$ ratios of RNase-treated 24h anlagen Hi-C vs. control 24h anlagen Hi-C across scaffolds > 5 Mb. Lower track is PCA1 value of RNase-treated 24h anlagen in 100 kb bins.
(B) Compartmentalization scores of RNase-treated 24h anlagen and control 24h anlagen.
(C) Frequency of different interaction types within RNase-treated 24h anlagen Hi-C and control 24h anlagen Hi-C.

| Timepoint | Samples | Total Reads | Total Uniquely Mapped Pairs | Pairs With ≥1 Mate Multimapping | Total Non-Duplicate Hi-C Contacts |
|---|---|---|---|---|---|
| Vegetative MIC | 2 | 524,216,869 | 100,191,043 | 332,005,536 | 73,405,616 |
| 24 hour anlagen | 2 | 571,744,886 | 252,885,143 | 199,574,586 | 178,108,763 |
| 48 hour anlagen | 1 | 103,915,592 | 58,626,828 | 38,127,701 | 39,863,562 |
| 24 hour anlagen +RNase | 1 | 101,044,731 | 57,640,945 | 35,313,022 | 38,088,780 |
| 24 hour anlagen RNase Control | 1 | 177,628,432 | 72,588,158 | 95,654,097 | 49,078,752 |

**Table S1. Summary of Hi-C datasets.**
